# Colorectal cancer-associated PCBP1 mutations disrupt protein stability in a dominant negative manner

**DOI:** 10.1101/2025.09.23.678070

**Authors:** Paige V. Blinkiewicz, Nicole C. Hoenes, Myra X. Afzal, Yihong Liu, Ulysses J. Hill, Berglind Bjarnadottir, T. Emme Burgin, Prerna Malaney

## Abstract

Mutations in RNA-binding proteins are increasingly identified in cancers through tumor sequencing and are correlated with disease progression, therapy response, and overall patient outcomes, underscoring the need to study them. Here, we focus on the RNA-binding protein Poly-C binding protein 1 (PCBP1), which binds target RNAs through K-homology (KH) domains to regulate RNA fate. PCBP1 is a tumor suppressor gene and hotspot missense mutations at leucine residues 100 and 102 are observed in colorectal cancer (CRC). PCBP1 mutations have been recurrently reported in CRC genome-wide mutation studies and are associated with poor clinical outcomes; however, their effects on PCBP1 expression and function remain largely unexplored. We show that cancer-associated mutations substituting leucine 100 and 102 with glutamine, proline, or arginine destabilize PCBP1, leading to increased protein turnover. The L100/L102 residues occur at the interface of the RNA-binding KH1 and KH2 domains, and our molecular dynamics simulations show that mutations at these residues disrupt the secondary structure of PCBP1. Additionally, these mutants display increased cytoplasmic localization. Importantly, mutant PCBP1 physically interacts with wild type PCBP1 and suppresses its expression through a dominant-negative mechanism. Together, our data demonstrate that CRC-associated PCBP1 mutations destabilize the protein and act as dominant-negative variants, revealing a novel mechanism of tumor suppressor inactivation in colorectal cancer.

## Introduction

RNA-binding proteins (RBPs) are key modulators of gene expression within the cell. They bind to RNA through conserved RNA-binding domains and modulate the stability, splicing, degradation, and translation of their target RNAs^1^. Given their control over global gene expression programs, aberrations in RBP expression or function result in pathogenic states. Increased accessibility to whole-exome sequencing has led to the identification of RBP mutations in both solid tumors and hematological malignancies^2,3^. These mutations are typically correlated with disease progression and response to therapy, ultimately affecting patient outcomes^3–9^ and making it critical to consider RBP mutations in the clinical management of malignancies.

Herein, we focus on an RBP called Poly-C binding protein 1 (PCBP1). *PCBP1* is an intronless gene that arose from the retro transposition of the fully processed mRNA of its homolog Poly-C binding protein 2 (*PCBP2*)^10^. Given its recent evolution, *PCBP1* is only found in mammals^10^. Despite its recent evolutionary emergence, *PCBP1* is an essential gene, and mice with a homozygous deletion of the *Pcbp1* allele are embryonic lethal^10,11^, indicating that it has acquired functions beyond those of PCBP2. Specifically, PCBP1 has a role in amino acid metabolism during development^12^ that likely renders it essential. Taken together, studies from mouse models indicate that both *PCBP1* and *PCBP2* have evolved to have non-redundant functions and are independently essential genes.

PCBP1 binds to poly-C rich sequences in mRNA through its RNA-binding KH domains and controls their stability, splicing, and translation^13^. While comprehensive mouse modeling for *PCBP1* in the context of tumor development is lacking, the gene is generally considered to be a tumor suppressor gene. Decreased PCBP1 protein expression is observed in lung cancer^14,15^, ovarian cancer^16^, and colon cancer^17^. Mechanistic studies have shown that PCBP1 suppresses oncogenic signaling pathways, including the Wnt/β-catenin^14^ and the MAPK pathway^18^, through post-transcriptional regulation of pathway components. In murine xenograft studies, reduced PCBP1 is associated with increased tumor growth and metastatic potential^18,19^. PCBP1 is also required for T-cell function and deletion of *Pcbp1* in murine T-cells causes blunting of antitumor immunity^20^. Together, these studies support a cell intrinsic and extrinsic tumor suppressive function for PCBP1.

In addition to changes in PCBP1 expression in tumors, mutations in *PCBP1* are observed in 15% of Burkitt lymphomas^21–23^ and 3-5% of colorectal cancers^24–28^. While mutations in *PCBP1* in Burkitt lymphoma are scattered along the entire length of the gene^21,22^, hotspot missense mutations in *PCBP1* are observed in colorectal cancers^29,30^. These mutations occur at leucine 100 and leucine 102 residues in the RNA-binding KH2 domain of PCBP1 and are associated with poor clinical outcomes in patients^31,32^. The *PCBP1* mutations in colorectal cancer generally co-occur with *APC* and *KRAS* mutations and are mutually exclusive with *TP53* mutations^33^. Despite the repeated identification of *PCBP1* mutations in colorectal tumor cohorts, the mechanism by which these mutations alter PCBP1 expression and function remains unclear.

Herein, we characterize the hotspot missense mutations that occur at residues L100/L102 in PCBP1 in colorectal cancer. The L100/L102 residues occur at the interface of the RNA-binding KH1 and KH2 domains, and mutations at these residues disrupt the secondary structure of PCBP1. Consequently, these mutations destabilized PCBP1 resulting in increased protein turnover. We observed that PCBP1 L100/L102 mutants can form hetero-oligomers with wild type PCBP1 and suppress its abundance levels in a concentration-dependent manner. Taken together, our data suggest that PCBP1 L100/L102 mutants act in a dominant negative manner to suppress wild type PCBP1 expression.

## Materials and Methods

### Analysis of Public Data

Data for PCBP1 mutations in colorectal cancer samples was accessed from cBioPortal on 20^th^ August 2025. Samples with the sample ID were treated as duplicates and only a single instance for these samples were considered for our analysis. Average values for variant allele frequencies were calculated for the duplicated samples and only considered once in our analysis. The TANGO webserver^34^ was used to determine the aggregation propensity of PCBP1. The webserver was accessed on 14^th^ June 2025 and the human PCBP1 sequence was input under the following parameters: N-term = N, C-term = N, pH = 7, Temperature = 298.15 K, and Ionic Strength = 0.02. For AlphaMissense predictions, data was accessed under the AlphaFold DB version 2022-11-01.

### Cell culture

HCT116 (ATCC #CCL-247) and HEC1A (ATCC #HTB-112) cells were cultured in McCoy’s 5A Media (Cytiva, SH30200.01). MC38 (Sigma Aldrich, SCC172) cells were cultured in DMEM (Dulbecco′s Modified Eagle′s Medium, Sigma, D5796). PC3 (ATCC #CRL-1435) and DU145 (ATCC #HTB-81) cells were cultured in RPMI (Cytiva, SH30027.01). All media was supplemented with 10% fetal bovine serum (FBS, Sigma, 12306C) and 1% Penicillin/Streptomycin (Sigma, P4333). PC3 were additionally supplemented with 1% HEPES (Thermo Scientific Chemicals, A1651622). The cells were maintained in a humidified incubator with 5% CO_2_ at 37°C.

### Plasmids

Human PCBP1, PCBP2, and hnRNPK point mutants were created using the nested PCR technique^35^. Q5 High Fidelity 2X Master Mix (New England Biolabs, M0492L) was used for PCR amplification. The gel purified products were digested and ligated into the pcDNA c-flag vector^36^ (Addgene plasmid #20011; http://n2t.net/addgene:20011; RRID: Addgene_20011) using XhoI and EcoRI sites. All plasmids contain a C-terminal Flag or HA tag, unless otherwise indicated as having an N-terminal tag. Sequences were verified using whole plasmid sequencing (Plasmidsaurus). Please see Supplemental Table 1 for primer sequences.

### Transfections

All plasmids were diluted to 250 ng/µl for transfections. For readout of Flag-tagged PCBP1 expression by western blot, cells were seeded in 6-well plates, allowed to reach 50% confluency, and then transfected with 2.5 µg of PCBP1, PCBP2, hnRNPK, or mutants thereof using JetPrime Reagent (Polyplus, 55-134). For western blot readout of dominant negative ability of PCBP1, PCBP2, or hnRNPK mutants, cells were seeded in 6-well plates and allowed to reach 50% confluency. They were then transfected with 200 ng of WT-HA-tagged plasmid with increasing amounts of WT-Flag-tagged plasmid (0, 75, 150, 300, 450, 600 ng) or MUT-Flag-tagged plasmid (0, 100, 200, 300, 400, 600, 800 ng). The total amount of plasmid DNA in each well was set to 1.0 µg, with the difference being made up using a pcDNA empty vector. For readout of co-immunoprecipitation, cells were seeded in 10cm dishes, allowed to reach 50% confluency, and transiently transfected with 4 µg PCBP1 WT plasmid and/or 5 µg PCBP1 L100Q plasmid. The amount of plasmid DNA in each well was set to 9.0 µg with pcDNA empty vector. For the readout of immunofluorescence, cells were seeded in 12-well plates, allowed to reach 50% confluency, and then transfected with 2.5 µg of Flag-tagged PCBP1 expression plasmids. Cells were harvested for western blots or processed for immunofluorescence experiments 48 hours post-transfection.

### Cell lysis and Western blotting

Cells were washed in PBS and lysed in either NP40 buffer (50mM Tris-HCl pH 8.0, 150mM NaCl, 1% NP40) or TN1 lysis buffer (50mM Tris-HCl, pH 8.0, 125 mM NaCl, 10 mM EDTA, 1% Triton-X 100, 16.5 mM Sodium Pyrophosphate, 10µM NaF) supplemented with Pierce protease and phosphatase inhibitors (Thermo Fisher Scientific, A32953 and A32957), unless otherwise stated. Lysates were cleared by centrifugation at 15,000 x g for 15 minutes at 4°C. In figure 5, cells were washed with PBS and lysed in denaturing Buffer (2% SDS, 2M Urea, 14% Sucrose, 1 mM NaF, 1 mM Sodium Orthovanadate, 25 mM β-Glycerophosphate) or 8M Urea supplemented with Pierce protease and phosphatase inhibitors. Whole cell lysates were then sonicated (continuous; 15% amplitude; 10 seconds on, 30 seconds off x 3 rounds). Following sonication, whole cell lysates were allowed to cool on ice, followed by centrifugation at 15,000 x g for 15 minutes at 4°C and further analyzed by western blot. Whole cell lysate protein concentration was determined using BCA Assay (Thermo Fisher Scientific, PI23227). Soluble proteins were boiled in 4X Laemmli buffer containing 10% β-mercaptoethanol, resolved on a 10% SDS-PAGE gel, and transferred to a Nitrocellulose or PVDF membrane by either wet or semi-dry transfer. Membranes were blocked for one hour at room temperature, probed with primary antibody overnight at 4°C, then probed with peroxidase conjugated secondary antibody for one hour at room temp. The antibodies used along with their conditions and dilutions are outlined in Supplemental Table 2. Protein bands were detected by chemiluminescence using a ChemiDoc MP Imaging System (BioRad). Western blots were quantified using BioRad Image Lab Software. Three replicates for each condition were analyzed using GraphPad Prism 10.4.1. p-values were determined using one-way ANOVA followed by Dunnett’s multiple comparisons test or a Mann-Whitney test.

### Molecular Dynamics Simulation

To try to develop a molecular-level rationale for the experimentally observed expression changes, molecular dynamics (MD) simulations for the fourteen variants plus wild-type were generated and analyzed. The PCBP1 KH2 domain structure remains to be determined; instead, the first conformer of the NMR structure for the KH1-KH2 domains of PCBP2^37^ (PDB ID: 2JZX) was used as the basis for modeling. This structure for PCBP2 is an identical homolog at the amino acid level to the corresponding region of PCBP1. Hydrogens were removed and added back at pH 6.5 using H++^38^. An N-methylamide group (NME) and an acetyl group (ACE) were manually added to the C- and ^39^N-termini of the PDB file, respectively, to prevent free-floating end charges. The LEaP program in the Amber22^40^ package was used to solvate the structure in water in an octahedral box with at least 10 Å between the protein and box edge, and to neutralize charge using Cl-counterions. The ff19SB force field and OPC3 water model^41^ were used. Hydrogen mass repartitioning (HMR) was performed using the parmed package in Amber22 to allow for time steps of 4 fs. All simulations were performed using the Amber22 package. The SHAKE algorithm was used to constrain the lengths of all hydrogen bonds unless otherwise stated. A cutoff distance for nonbonded interactions was set to 8 Å. First, the structure was minimized without SHAKE over 2000 steps (1000 steps of steepest descent followed by 1000 steps of conjugate gradient). The system was then heated from 0 to 310K over 10,000 steps using a 2 fs timestep at constant volume. During heating, temperature was controlled using a Langevin thermostat^42^ with a collision frequency of 2.0 ps^-1^. Following heating, a short 2 ns equilibration was carried out under constant pressure (1 bar, 310 K) with a 4 fs timestep. Mutations were introduced to the equilibrated wild-type (WT) structure with PyRosetta^43^. The variants and wild type then underwent minimization and heating as described previously, followed by a 100 ns production simulation. Production was performed under the same conditions as the previous 2 ns equilibration. For equilibration and production, temperature was held at 310 K using an Andersen thermostat^44^ with a velocity randomization every 4 ps. Trajectory outputs were saved every 1000 steps. Six independent trajectories were generated per sequence, each with random starting velocities.

### MD Trajectory Analysis

Following the generation of the MD simulations, the production trajectories were analyzed for relative conformational flexibility between protein domains (e.g., how much one domain moves relative to the other). To do this, the PCBP2 model was separated into two domains: residues 10-88 (KH1 domain) and 98-169 (KH2 domain) based on the residue indices from the original PDB structure. Residues 89 to 97 are located on a flexible loop region and were not included in this calculation. To calculate the relative motion of the KH1 domain to the KH2 domain, the KH2 domain was fixed in space while the KH1 domain was overlaid on top of the first frame of each simulation using the superpose function from the Pytraj library^45^. The root mean square fluctuation (RMSF) for each residue of the KH1 domain was then calculated using the Pytraj function rmsf. The RMSF values for the KH1 domain were averaged across residues, and these residue-average values were then averaged across the six replicate trajectories. The calculated average RMSF values represent the relative motion or displacement of the KH1 domain relative to the KH2 domain for each variant. The average RMSFs in angstroms were plotted against the experimental protein fold expression values to determine whether a relationship was present.

### Immunofluorescence

Immunofluorescence assays were conducted as previously described by Deng et al., 2023^46^. Briefly, HCT116 cells were seeded onto standard 18 mm glass coverslips (Electron Microscopy Sciences, 7222201) and transfected as described above. Post transfection, cells on the coverslips were fixed with 3.5% paraformaldehyde. After fixation, the cells were permeabilized and blocked using Triton X-100 and 2% BSA. The cells were then incubated with primary antibodies, washed, and subsequently treated with fluorescent secondary antibodies and DAPI. Finally, the samples were washed again and mounted onto glass slides using ProLong Gold antifade (Invitrogen, P36934). Representative images are shown of three replicates performed. The antibodies used along with their condition and dilutions are outlined in Supplemental Table 2. Samples were imaged using a Ti2-E Inverted Microscope and analyzed using Image J and NIS Elements software.

### Droplet digital PCR

Droplet digital PCR was performed to determine the copy number of *PCBP1* in HCT116 cells. The standard BioRad QX100 Droplet Digital PCR protocol was followed. We used two reference genes *PPIA* and *POL2RA*. We calculated PCBP1 copy number using this formula: PCBP1 CNV = (PCBP1 events / reference gene events) × 2 (reference gene CNV).

### Generation of PCBP1^L100Q/WT^ and PCBP1^+/-^ cell lines

For generation of the PCBP1 ^L100Q/WT^ cells, HCT116 cells were obtained from ATCC (#CCL-247) and subjected to CRISPR/Cas9 transfection with sgRNA (5’ - GGCCGGCACCACCAGCCTCA - 3’) targeting the nucleotides around position 300 for generation of the PCBP1^L100Q/WT^ cell line. Through homology directed repair (HDR) (donor sequence: 5’ - CGCCTTTCCCAATCAGGGAGCCGCACTGGGTGGCCGGCACCACGAGTCTCTGGGTGACCGGGGGCCTGCTGGCCGCGGTACTGTTGGTCATGGAGCTGTT - 3’), a leucine to glutamine mutation was introduced at amino acid position 100 in one allele of PCBP1. The presence of the mutation was confirmed by sequencing (non allele specific PCBP1-F 5’ - GACTTGACCACGTAACGAGC - 3’ and on allele specific PCBP1-R 5’ - GCGGAGAAATGGTGTGTTGT - 3’). Sequencing was also performed showing that PCBP2 remained wild type post editing. Four clones harboring the PCBP1^L100Q/WT^ mutation were selected for downstream experiments (clones 9, 16, 177, and 320). For generation of PCBP1^+/-^ cells, HCT116 cells were subjected to CRISPR/Cas9 transfection with sgRNA (5’ - UCCGUGCAUAAGAAGCCGAA - 3’) targeting the nucleotides around the start site for generation of the PCBP1^+/-^ cells. The presence of the knocked-out allele was confirmed by sequencing (PCBP1-F 5’ - GCAGTTTTGGGCCTACACCT - 3’ and PCBP1-R 5’ - CGTACTCTCGCGGATCTCTT - 3’). Four clones were selected for downstream experiments (clones 1, 17, 19, and 33).

### Cellular Thermal Shift Assay

Cellular thermal shift assay was performed as described in Malaney et al^47^. Briefly, HCT116 PCBP1^WT/WT^ and HCT116 PCBP1^L100Q/WT^ cells were plated into T-175 flasks and allowed to reach 80% confluence. On day of collection, cells were trypsinized and spun down at 300 x g for 5 min and then washed with 1X PBS. The cell pellet was reconstituted in 1X PBS and cell counting was performed. The cells were spun down at 300 x g for 5 min and supernatant was aspirated off pellet. The pellet was resuspended in 1X PBS with protease inhibitors to a concentration of 30 million cells/ 1 mL PBS and 100ul of cell suspension were aliquoted into eight 0.2 mL tubes. One tube for each cell line was kept as the room temperature (RT) sample, while the remaining tubes were placed in a thermocycler for 8 minutes with a temperature gradient increasing in 2°C increments from 42°C to 52°C. The tubes were then snap-frozen in liquid nitrogen, followed by a quick thaw at 25°C. The freeze/thaw process was repeated three times. Lastly, the lysate was transferred to a 1.5 mL microcentrifuge tube, centrifuged at 20,000 x g for 20 minutes at 4°C, and the resulting supernatant was boiled with 4X Laemmli buffer containing 10% β-mercaptoethanol. Equal volumes of lysate were used in western blots.

### Protein Degradation Assays

HCT116 PCBP1^WT/WT^ and HCT116 PCBP1^L100Q/WT^ cells were seeded in 6-well plates and allowed to reach 70-80% confluency. They were then treated with cycloheximide (50 µM) for 0, 9, and 18 hours. Upon collection, cells were washed in 1X ice cold PBS and lysed in NP40 or TN1 lysis buffer supplemented with Pierce protease and phosphatase inhibitors (Thermo Fisher Scientific, A32953 and A32957). Lysates were cleared by centrifugation at 15,000 x g for 15 minutes at 4°C and analyzed by western blot. To determine the pathways through which protein degradation was occurring, cells were co-treated with cycloheximide and either a lysosome inhibitor Bafilomycin A1 (100nM, MedChemExpresss, HY-100558) or proteasome inhibitors MG132 (10 µM, Cayman chemical, 1001262810) or Bortezomib (30 nM, MedChemExpress, HY-10227).

### Protein Degradation Assay Quantification

We utilized MATLAB and its integrated development environment (IDE) to create custom software to analyze the data from the cycloheximide chase assays. First, the western blot image is imported, converted to grayscale, and a Gaussian blur is applied to remove noise. The user is then prompted to select regions of interest (ROIs), which should encompass each individual band. For each ROI, canny edge detection is employed to determine the band’s edges, followed by morphological closing. An image displaying the bands as detected by Canny is generated, with the bands highlighted in green. It is crucial to manually verify the accuracy of the Canny detection at this stage, as it can be sensitive to ROI selection. If the bands are not adequately detected, the ROI should be re-selected. For bands exhibiting clear decay, calculated half-life values were highly reproducible, with an average variation of less than two hours between runs. In contrast, for bands displaying little or no change in intensity over the time course, half-life estimates were more variable. This behavior is expected and does not affect the reliability of the analysis for bands where meaningful decay is present. Normalization is then performed. This process sets the first concentration for each cell line to one, with the remaining values generally being less than one. When little to no measurable decay is observed, the program will output a half-life of infinity, reflecting the absence of decay in the control sample. Such runs are excluded from average half-life calculations for experimental groups. The code then fits exponential decay curves (1*exp(-b*x)) to data for multiple replicates, computes key metrics, and plots the results. Finally, it should be noted that the specific settings, such as the canny edge detection threshold and the level of morphological closing, are optimized for this particular set of images and may not be suitable for all images.

### RNA Isolation and RT-qPCR Analysis

HCT116 PCBP1^WT/WT^ and HCT116 PCBP1^L100Q/WT^ cells were seeded in 6-well plates and allowed to reach 80% confluency. Cells were washed with 1X PBS followed by RNA isolation using the Zymo Quick-RNA MiniPrep kit (Zymo Research, R1055) according to the manufacturer’s protocol. 500 ng of isolated RNA was converted to cDNA using the iScript Reverse Transcription Supermix kit (BioRad, 1708841BUN) according to the manufacturer’s protocol. Quantitative PCR was performed for *PCBP1* and *PPIA* transcript using the iTaq Universal SYBR Green Supermix (BioRad,1725124) according to the manufacture’s protocol and run on a QuantStudio 3 (Thermo Fisher Scientific). The primers used are outlined in Supplemental Table 3. Data are the average of three biological replicates, performed in triplicate. Data analysis was performed using the ΔCq^48^ method and significance was determined using a one-way ANOVA with post-hoc Dunnett’s test in GraphPad Prism.

### Actinomycin Chase Assay

HCT116 PCBP1^WT/WT^ and HCT116 PCBP1^L100Q/WT^ cells were seeded in 6-well plates and allowed to reach 60-70% confluency. They were then treated with actinomycin D (5 mg/ml, Sigma, A9415) for 0, 1, 2, 4, 6, and 8 hours followed by RNA isolation and qPCR as described above. Data are the average of three biological replicates, performed in triplicate. Data analysis was performed using the ΔCq method^48^ and significance was determined using a two-way ANOVA with post-hoc Dunnett’s test in GraphPad Prism.

### Co-Immunoprecipitation

Whole cell lysates were prepared using Pierce IP Lysis Buffer (25 mM Tris HCl pH 7.4, 150 mM NaCl, 1% NP-40, 1 mM EDTA, 5% glycerol) supplemented with phosphatase and protease inhibitors (Thermo Fisher Scientific, A32953 and A32957). Protein concentration was determined by BCA Assay (Thermo Fisher Scientific, 23223). For immunoprecipitation assays, 300 µg of protein was incubated with HA beads (Thermo Fisher Scientific, 88836) or Flag beads (Sigma, M8823) for 2 hours on a rotator at room temp. The resulting bead/immuno-complex was then washed 3 times in 500 ul of TBST and boiled in 100 µl of 2X SDS Laemmli buffer containing 10% β-mercaptoethanol. Co-immunoprecipitated proteins were subsequently analyzed by western blot. Western blots are representative of three independent experiments.

## Results

### Hotspot mutations occur in PCBP1 in colorectal cancer

We queried the cBioPortal database to identify mutations that occur in PCBP1 in colorectal cancer. Of the 77 reported PCBP1 mutations in colorectal cancers, we observe that 66 (85%) occur at residues leucine 100 and leucine 102 representing a hotspot for mutations (Figure 1A). Upon surveying the different PCBP1 mutations, we observe that leucine 100 and 102 are mutated to residues glutamine, proline, or arginine (Q, P, or R), with the leucine to glutamine change at residue 100 being the most commonly observed mutation (Figure 1B, herein abbreviated as L100Q). While there are mutations that occur in residues other than leucine 100/102, these are relatively uncommon (Figure 1B). The variant allele frequency for L100/L102 mutations lies between 0.3 to 0.5 (Figure 1C) indicating that PCBP1 wild type and mutant alleles co-exist in tumor samples. Given that PCBP1 is an essential gene, it is no surprise that we do not see variant allele frequencies of 1. These observed variant allele frequencies are in line with those reported for other RNA-binding proteins mutated in cancers such as SRSF2^49^, U2AF1^50^, and SF3B1^49^.

**Figure 1:**
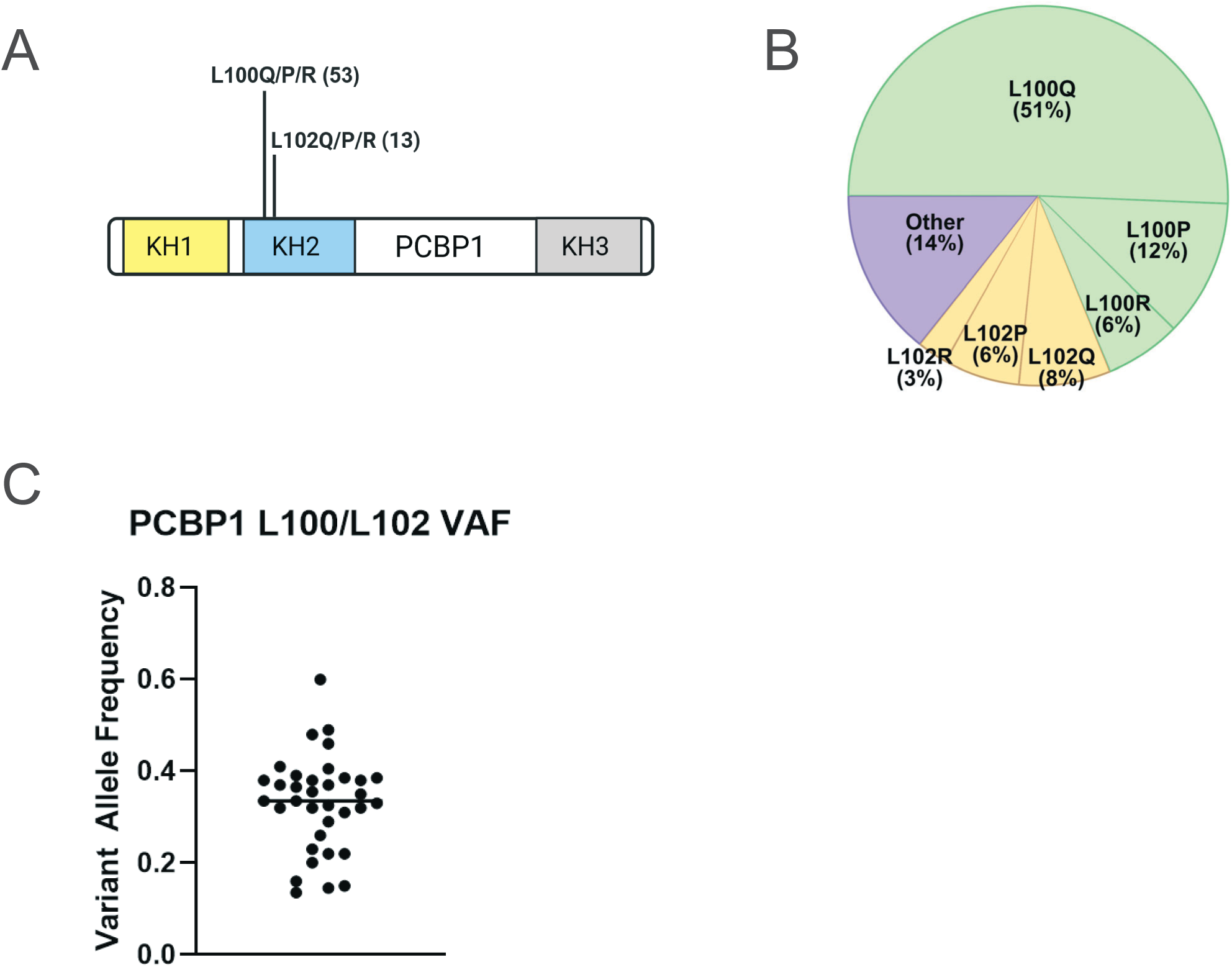
PCBP1 hotspot mutations in colorectal cancer. **(A)** Diagram depicting the hotspot mutations that occur in PCBP1 in colorectal cancer. **(B)** Proportion of different PCBP1 mutations observed in colorectal cancer. **(C)** Variant allele frequency for mutant PCBP1 in human colorectal cancer patient samples (n=35). All mutation data has been obtained from cBioPortal.

### Mutations at L100/L102 disrupt PCBP1 protein expression and conformation

While the structure of the PCBP1 KH2 domain remains to be determined, an NMR structure for the KH1-KH2 domains of PCBP2 is reported (PDB ID: 2JZX)^37^ which are 94% identical at the amino acid level to PCBP1. The KH domains of PCBP2 have a classical type-I KH domain fold which consists of three α-helices and three β-strands arranged in the order β1-α1-α2-β2-β3-α3^37^. The KH1 and KH2 domains of PCBP2 form an intramolecular pseudodimer, and the dimerization interface is comprised of strand β1 and helix α3 of both the KH1 and KH2 domains^37^. The KH1-KH2 dimerization interface is primarily hydrophobic in nature. The L100/L102 residues are contained within the β1 strand of the KH2 domain and are located at the dimerization interface between the KH1 and KH2 domains^29,37^ (Figure 2A). Given that the KH1/KH2 interaction stabilizes the PCBP protein^37^, and studies show propensity for mutations at inter-domain contacts to impact protein stability^51^, we wanted to determine the protein expression levels of the cancer-associated PCBP1 mutants. To this end, we cloned all cancer-associated PCBP1 L100/L102 mutations (L100Q/P/R and L102Q/P/R) into a mammalian expression vector with a C-terminal Flag tag and transiently expressed them in the human colorectal cancer cell line, HCT116. We observed that all PCBP1 L100/L102 mutants have lower expression levels compared to their Flag-tagged wild type counterparts (Figure 2B). HA-tagged wild type PCBP1 was included in the experiments as an additional control with a different tagging epitope.

**Figure 2:**
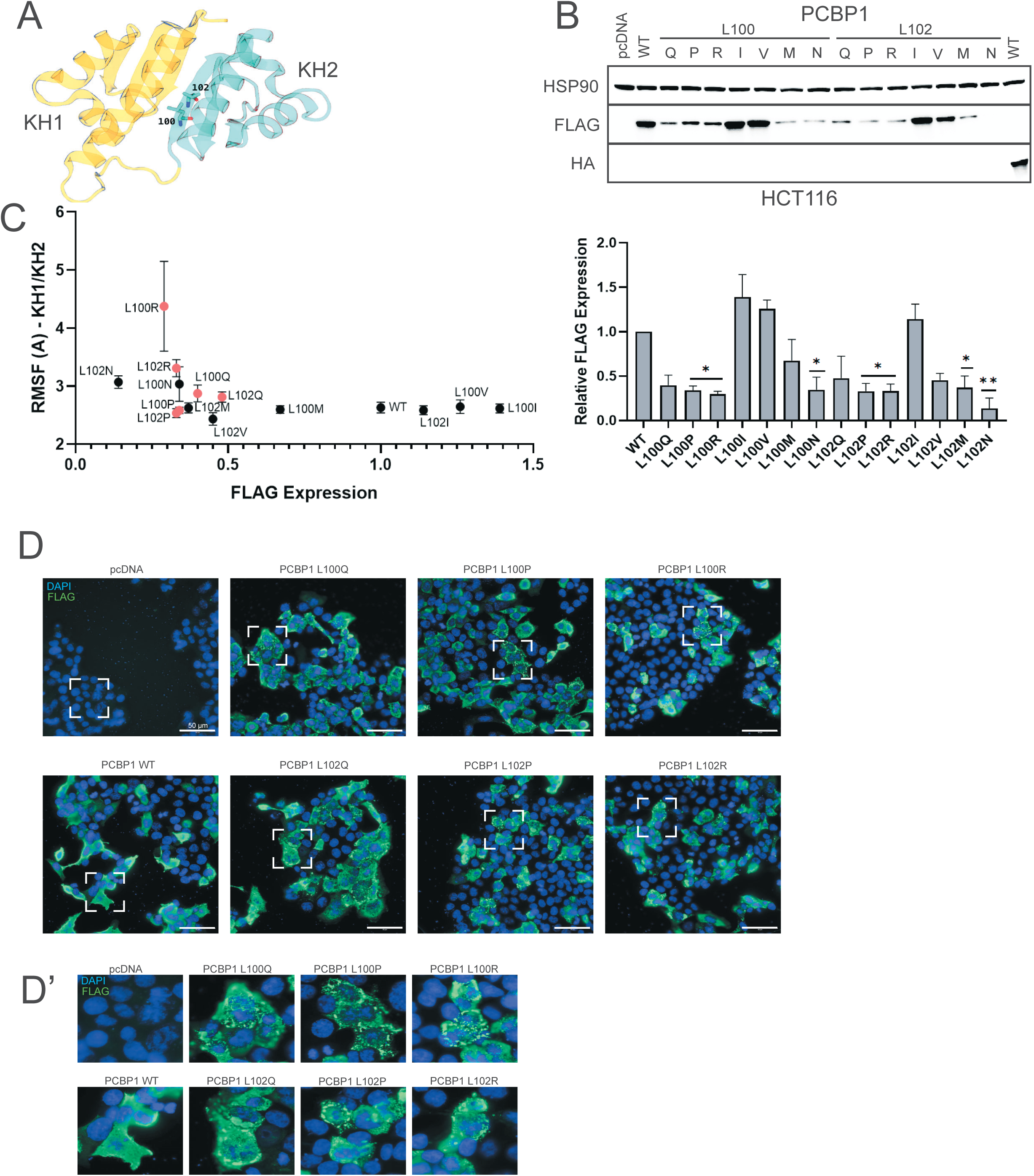
Mutations at L100/L102 disrupt PCBP1 protein expression and conformation. **(A)** Post-equilibration wild-type structure for KH1-KH2 domain structure from PCBP2 (PDB ID: 2JZX). The L100 and L102 residues are indicated. **(B)** Western blot showing expression level for C-terminal Flag-tagged PCBP1 plasmids upon transient transfection in HCT116 cells. Quantification of the western blot was done using three independent replicates and p-values were calculated using one-way ANOVA with a post hoc Dunnett’s Test (* p<0.05; ** p<0.01). **(C)** Relative root mean square fluctuation of the KH1-KH2 domains correlated with the expression levels of the various PCBP1 mutants in HCT116 cells. RMSF across residues in the KH1 domain are calculated relative to residues in the KH2 domain. **(D and D’)** Sub-cellular localization of PCBP1 mutants, as assessed by Flag immunofluorescence, upon transient transfection in HCT116 cells. Representative image is shown (scale=50 µm; n=3).

For all cancer-associated PCBP1 L100/102 mutations (Figure 1B), we observed that leucine was replaced with larger, charged, or structure-breaking amino acids. To understand whether the charge or size of the mutant amino acid contributes to decreased expression, we created a series of rationally designed mutants PCBP1 L100I/V/M/N and L102I/V/M/N. Substitution of L100/L102 with an isomer, isoleucine, restored stability to wild type levels. Similar results were observed with a valine substitution. Replacement with methionine, a non-polar but bulkier residue, resulted in a slight decrease in PCBP1 stability. However, replacement with a polar residue, asparagine, dramatically reduced PCBP1 stability (Figure 2B). Together, these findings indicate that a combination of charge and size of the mutated amino acids disrupt the hydrophobic nature of the KH1-KH2 domain interface, which may contribute to decreased stability of cancer-associated PCBP1 mutations. These observations were recapitulated in a murine colorectal cancer cell line, MC38 (Supplementary Figure 1A). We also tested N-terminal Flag-tagged PCBP1 mutants (L100Q and L100I) in HCT116 cells and observed a similar decrease in PCBP1 expression for the L100Q mutant, indicating that this loss of stability is not due to epitope tagging (Supplementary Figure 1B).

We then used AlphaMissense^52^ to predict the pathogenicity of mutations in PCBP1. The results for residues L100 and L102 are highlighted in Supplementary Figure 1C. Here, we observe that all of the colorectal cancer-associated PCBP1 mutations (L100Q/P/R and L102Q/P/R) are likely pathogenic. The rationally designed PCBP1 L100N and L102N mutations are also predicted to be pathogenic. All of these putative pathogenic mutants show reduced expression, matching our data from the colorectal cancer cell lines HCT116 and MC38 (Figure 2B, Supplementary Figure 1A). The rationally designed PCBP1 L100I/V/M and L102I/V/M mutations are predicted by AlphaMissense as benign, and consistently these mutants have comparable expression to wild type PCBP1 in our data (Figure 2B, Supplementary Figure 1A).

The KH1 and KH2 domains in PCBP2, and therefore by extension likely in PCBP1, form a stable intra-molecular dimer with the L100/L102 residues at the dimerization interface^37^. We therefore wanted to interrogate whether the PCBP1 mutations impact the dimerization interface and whether this is correlated with protein expression levels. Using the PCBP2 KH1-KH2 domain structure (PDB ID: 2JZX), we used molecular dynamics simulations to measure the flexibility and movement of the KH1 and KH2 domains, relative to each other, for wild type and L100/L102 mutants. Here, we observed an inverse relationship between KH1-KH2 inter-domain flexibility and PCBP1 protein expression levels in HCT116 cells (Figure 2C). Wild type PCBP1 that has high expression levels shows relatively low inter-domain movement. However, some of the cancer-associated PCBP1 L100/L102 mutants show much higher inter-domain flexibility (Figure 2C). In contrast, the rationally designed PCBP1 L100V, L100I, and L102I mutants have comparable inter-domain flexibility to wild type PCBP1. The two mutants PCBP1 L100P and L102P have very low expression yet seem to have low inter-domain flexibility comparable to wild type PCBP1. It is likely that these highly destabilized mutants are unable to fold properly in an experimental setting; however, the models do not take initial folding into account and assume the protein is already folded, which might explain the deviation to the otherwise strong trend between low inter-domain flexibility and high expression. We observe similar results with all of our PCBP1 mutants in MC38 cells (Supplementary Figure 1D). Together, our results indicate that the cancer-associated PCBP1 mutations likely disrupt the association between the KH1-KH2 domains resulting in impaired protein stability.

A recent study demonstrated that missense mutations that impact protein stability result in protein mislocalization^53^. To determine whether the L100/L102 mutations also alter PCBP1 sub-cellular localization, we performed immunofluorescence experiments in HCT116 cell lines transfected with Flag-tagged PCBP1 variants. Our data indicate that while wild type PCBP1 shows diffuse cellular staining^54^, cancer-associated PCBP1 L100/L102 mutants mislocalize in cytoplasmic punctate structures (Figure 2D and D’). These observations are in line with those reported by Porter et al^29^. This mislocalization is observed for the PCBP1 L100N and L102N mutants but not the PCBP1 L100I/V/M and L102I/V/M mutants (Supplementary Figure 1E and E’).

### Mutations at L100/L102 result in reduced expression of other KH domain containing RNA-binding proteins

Given that the cancer-associated PCBP1 mutations occur in the RNA-binding KH2 domain of PCBP1, we sought to determine whether these L100/L102 residues are conserved in other KH-domain containing RNA-binding proteins. We specifically focused on the most ubiquitously expressed KH-domain containing RBPs in mammalian cells, PCBP2 and hnRNPK^55^. We observed that the L100/L102 residue is conserved in PCBP2 and hnRNPK (Figure 3A). Upon mutating the L100/L102 residues in PCBP2 to Q/P/R, we observed a similar decrease in expression in HCT116 (Figure 3B) and MC38 cells (Supplementary Figure 2A). The corresponding residues L147/L149 in hnRNPK, when mutated, also result in lowered expression levels in HCT116 (Figure 3C) and MC38 cells (Supplementary Figure 2B). We observe that the reduction in expression is more profound for hnRNPK compared to PCBP2. The rationally designed hnRNPK L147I/V/M and L149I/V/M mutants have restored protein expression levels, whereas the L147N and L149N mutants have reduced expression levels. The hnRNPK L147M and L149M mutants are more stable than their PCBP1 counterparts, indicating that for hnRNPK it is likely the charge of the substituted amino acid that results in compromised protein expression (Figures 2B, 3C, Supplementary Figures 1A, 2B).

**Figure 3:**
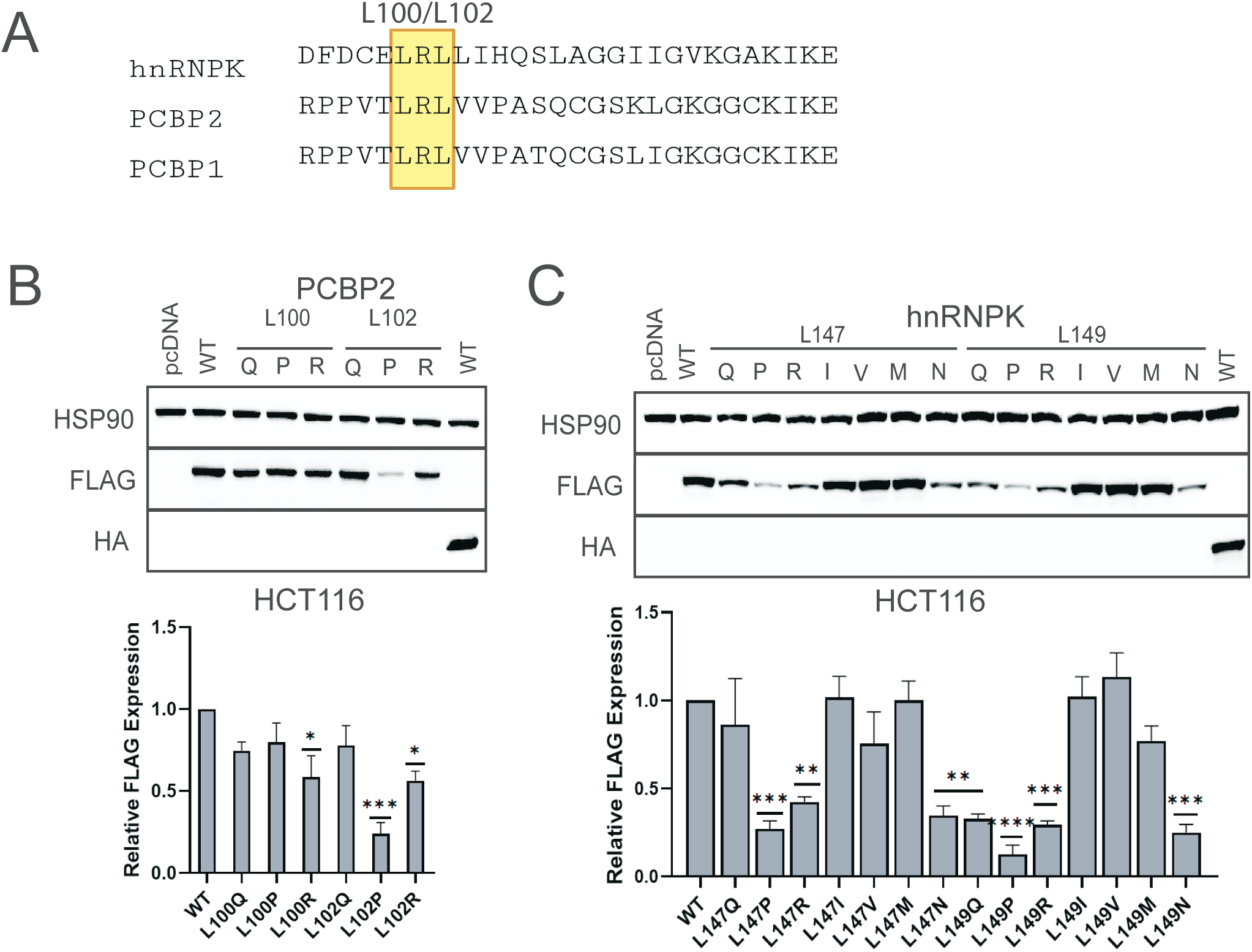
Mutations at L100/L102 result in reduced expression of other KH domain containing RNA-binding proteins. **(A)** Amino acid sequence alignment for hnRNPK, PCBP2, and PCBP1 around leucine residues 100 and 102. **(B & C)** Western blot showing expression level for C-terminal Flag-tagged PCBP2 and Flag-tagged hnRNPK plasmids respectively upon transient transfection in HCT116 cells. Quantification of the western blot was done using three independent replicates and p-values were calculated using one-way ANOVA with a post hoc Dunnett’s Test (* p<0.05; ** p<0.01; *** p<0.001; **** p<0.0001).

We then interrogated the COSMIC cancer mutation database to identify whether L100/L102 residue mutations were observed in PCBP2 or hnRNPK in cancer. We identified that PCBP2 L100R mutations occur in prostate cancer and hnRNPK L149M mutations occur in endometrial cancers. We transiently transfected Flag-tagged PCBP2 L100R plasmids into prostate cancer cell lines PC3 and DU145 and found reduced expression of the mutant (Supplementary Figure 2C-D), which is consistent with our results from HCT116 and MC38 cell lines. We transiently transfected Flag-tagged hnRNPK L149M plasmids into an endometrial adenocarcinoma cell line, HEC1A, and observed minimal reduction in expression levels compared to hnRNPK WT, which is also consistent with our results from HCT116 and MC38 cell lines (Supplementary Figure 2E).

### The L100Q mutation destabilizes PCBP1 protein resulting in increased protein turnover

To verify that the effects of PCBP1 L100/L102 mutations on expression levels were not an artifact of transient transfections, and in order to study cancer-associated PCBP1 mutations in a more disease-relevant model, we used CRISPR-editing to create HCT116 PCBP1 L100Q mutant cell lines. We selected the L100Q mutation as it is the most commonly observed mutation in human patient samples (Figure 1B). We created four clones of these cell lines: clones 9, 16, 177 and 320 (Figure 4A). HCT116 cells have two PCBP1 alleles (Supplementary Figure 3A), and our mutant clones have one PCBP1 wild type and one L100Q allele (HCT116 PCBP1^L100Q/WT^). We screened 800 clones and were unable to create cells with a homozygous PCBP1 L100Q mutation, likely due to the essentiality of the PCBP1 gene. We tested total PCBP1 protein levels in these CRISPR-edited cell lines and found lower total PCBP1 levels in our clones (Figure 4B).

**Figure 4:**
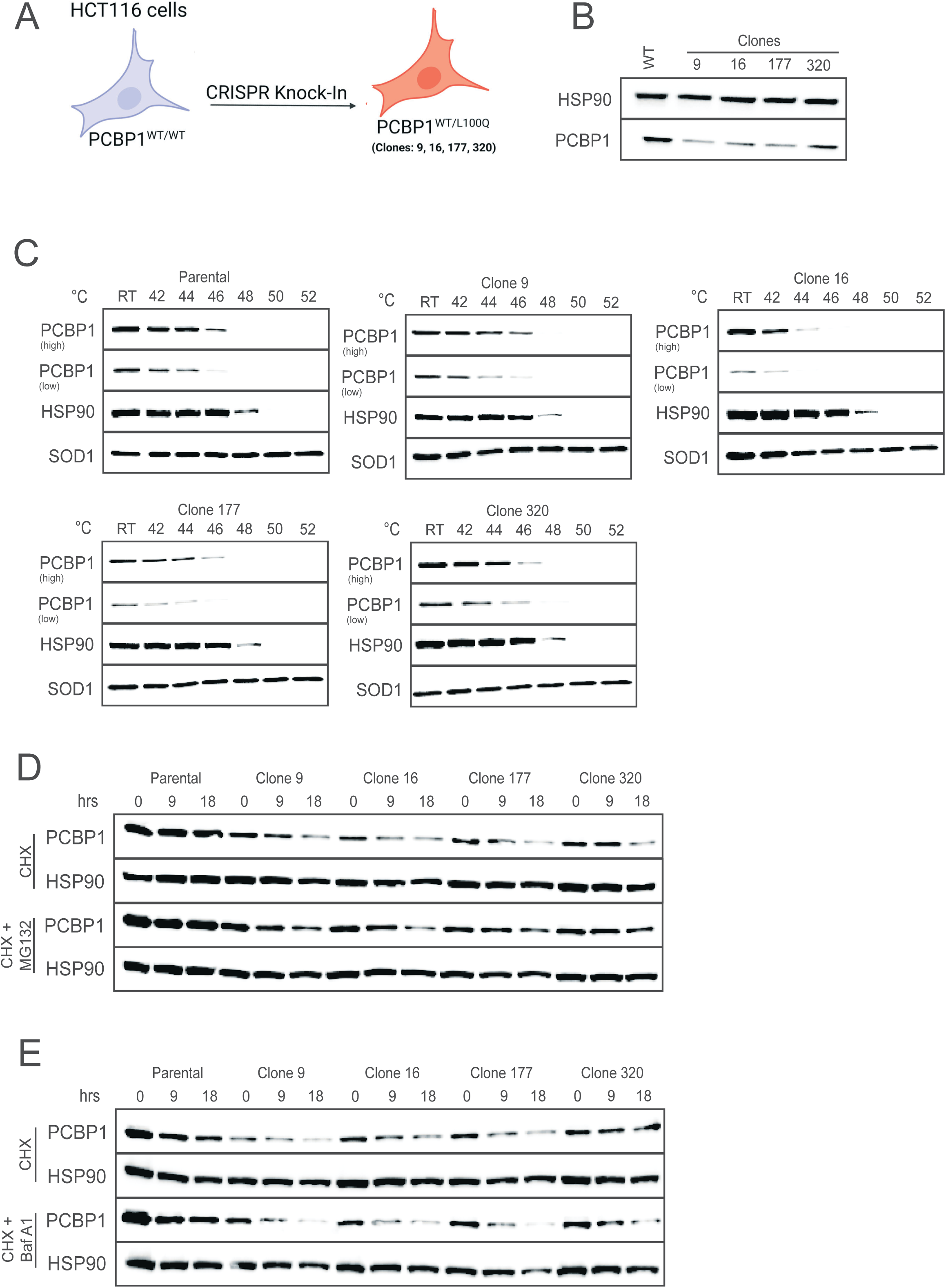
The L100Q mutation destabilizes PCBP1 protein resulting in increased protein turnover. **(A)** Generation of CRISPR edited HCT116 PCBP1^L100Q/WT^ cell lines. **(B)** Western blot showing expression level for total PCBP1 in HCT116 PCBP1^WT/WT^ and HCT116 PCBP1^L100Q/WT^ cells. **(C)** Cellular thermal shift assay to determine the melting temperature of total PCBP1 in HCT116 PCBP1^WT/WT^ parental cells and HCT116 PCBP1^L100Q/WT^ mutant clones. HSP90: positive control; SOD1: negative control. Representative image is shown (n=3). **(D)** Cycloheximide chase assays with and without proteasomal inhibitor, MG132, in HCT116 PCBP1^WT/WT^ and HCT116 PCBP1^L100Q/WT^ cells. Representative image is shown (n=3). **(E)** Cycloheximide chase assays with and without lysosomal inhibitor, bafilomycin A1, in HCT116 PCBP1^WT/WT^ and HCT116 PCBP1^L100Q/WT^ cells. Representative image is shown (n=3).

Next, we wanted to understand the mechanistic basis for the reduced expression of PCBP1 L100Q. Total RNA levels of *PCBP1* were largely unaltered or slightly increased in the HCT116 mutant cell lines indicating that this is likely not a transcriptional mechanism (Supplementary Figure 3B). We then assayed *PCBP1* mRNA stability using actinomycin D chase assays and observed no change in PCBP1 transcript stability in the PCBP1^L100Q/WT^ cells (Supplementary Figure 3C). Together, these experiments indicate that PCBP1 L100Q is likely destabilized at the protein level. We therefore performed a Cellular Thermal Shift Assay (CETSA) in our parental HCT116 PCBP1^WT/WT^ and PCBP1^L100Q/WT^ cells and observed a lower melting temperature for PCBP1 in the mutant clones (Figure 3C). HSP90, which melts at ∼46°C^56^, was included as a positive control and SOD1, which melts at 95°C^57^, was included as a negative control in this experiment. We also performed cycloheximide chase assays and established that PCBP1 has a shorter half-life in the mutant clones (half-life >18 hours for parental cells, 12.309 ± 1.586 hours for clone 9, 10.534 ± 1.357 hours for clone 16, 11.267 ± 2.158 hours for clone 177, and 15.001 ± 2.598 hours for clone 320, Figure 4D and Supplementary Figure 3D).

Having established that PCBP1 L100Q is destabilized at the protein level, we wanted to investigate whether we could rescue this destabilization using protein degradation inhibitors. We first used MG132, a proteasome inhibitor, and observed a partial rescue of PCBP1 expression levels in our mutant cell lines (Figure 4D and Supplementary Figure 3E). The half-lives for PCBP1 increased to 18.355 ± 1.762 hours, 12.451 ± 1.924 hours, 15.898 ± 3.427 hours and 22.672 ± 6.048 hours for clones 9, 16, 177 and 320 respectively. Similar results were observed with bortezomib, another proteasome inhibitor (Supplementary Figure 3F). Interestingly, no rescue of PCBP1 expression was observed with a lysosomal inhibitor, bafilomycin A1 (Figure 4E), indicating that PCBP1 mutants are likely not turned over through the lysosome. Together, these findings show that PCBP1 L100Q is destabilized, and its expression can be rescued in part through the blockade of protein degradation pathways.

### PCBP1 L100/L102 mutant proteins are insoluble

Our molecular dynamics simulation data suggests disrupted inter-domain interactions between the KH1 and KH2 domains for PCBP1 mutants. Given that changes in secondary and tertiary structure can cause protein aggregation, we assessed the aggregation potential of PCBP1. Using an aggregation prediction software, TANGO, we observe that the amino acid residues flanking the L100/L102 amino acids, F72-I76 and I162-167, have high aggregation potential (Figure 5A). For protein isolation, we use mild buffers containing non-ionic detergents like NP40 and Triton X, and using these buffers, we observe lower PCBP1 expression levels for the L100/L102 mutants. To interrogate whether the PCBP1 L100/L102 mutants are insoluble, we used harsher chaotropic protein isolation buffers to solubilize them. Using a denaturing buffer (containing 2% SDS and 2M urea), we recovered higher amounts of the Flag-tagged PCBP1 mutants in HCT116 cells (Figure 5B). We also used the denaturing buffer to lyse our HCT116 PCBP1^L100Q/WT^ clonal cell lines and found that we could recover higher amounts of PCBP1 protein in the PCBP1^L100Q/WT^ cells (Figure 5C). As a negative control, we included HCT116 PCBP1^+/-^ cells (clones 1, 17, 19 and 33) wherein we observed no such recovery of PCBP1 expression. We also repeated these experiments by lysing cells in 8M Urea and observed similar results (Figure 5D-E). Together, these data indicate that a fraction of the PCBP1 L100/L102 mutants are insoluble and likely aggregation-prone, which contributes to their lower recovery levels in the soluble fraction.

**Figure 5:**
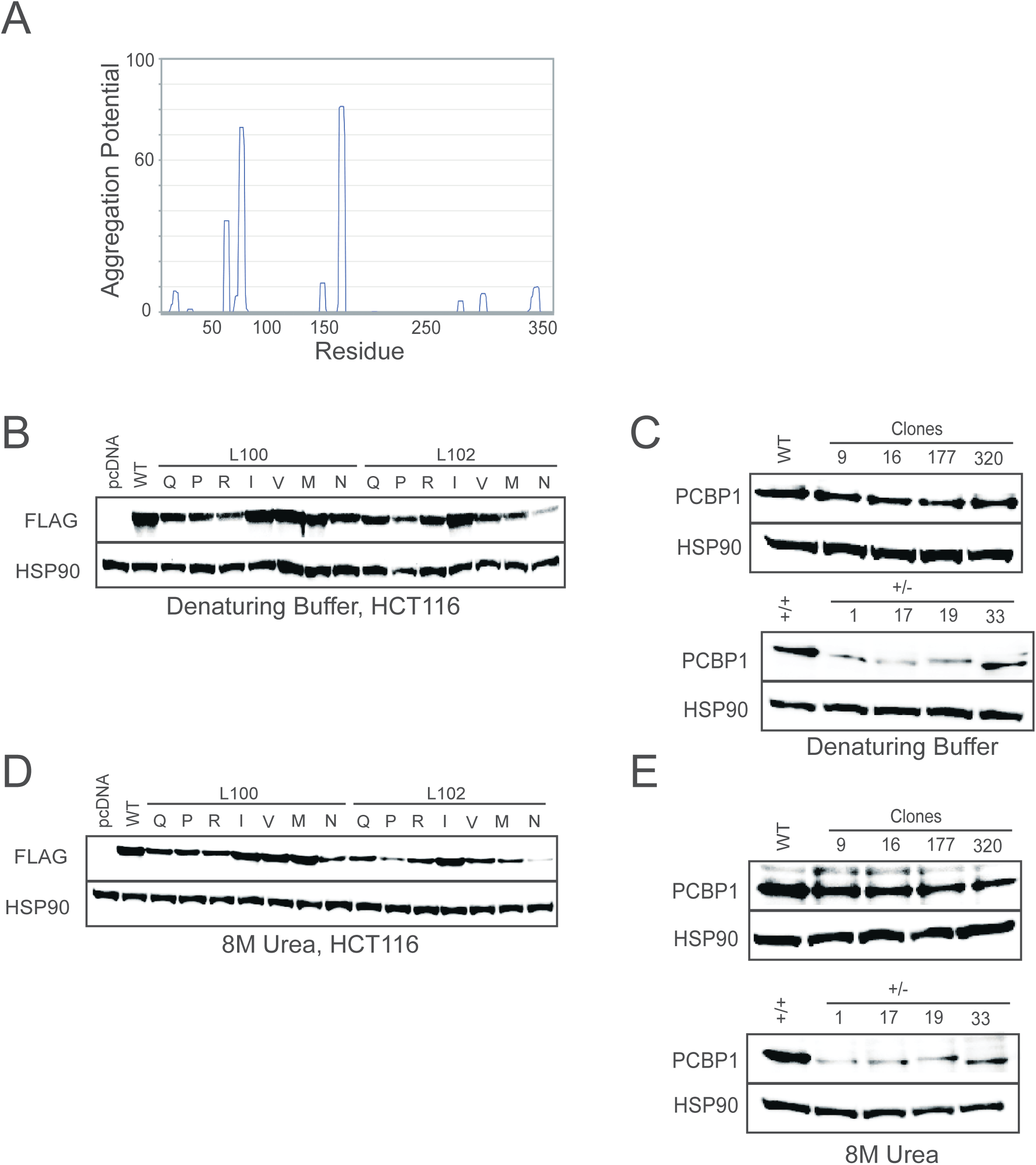
PCBP1 L100/L102 mutant proteins are insoluble. **(A)** TANGO analysis of the PCBP1 wild type amino acid sequence to determine aggregation potential. **(B)** Western blot showing expression level for C-terminal Flag-tagged PCBP1 plasmids upon transient transfection in HCT116 cells that were lysed in denaturing buffer. Representative image is shown (n=3) **(C)** Western blot showing expression level for total PCBP1 in HCT116 PCBP1^WT/WT^, HCT116 PCBP1^L100Q/WT^ cells, HCT116 PCBP1^+/+^ and HCT116 PCBP1^+/-^ that were lysed in denaturing buffer. Representative image is shown (n=3). **(D)** Western blot showing expression level for C-terminal Flag-tagged PCBP1 plasmids upon transient transfection in HCT116 cells that were lysed in 8M urea. Representative image is shown (n=3) **(E)** Western blot showing expression level for total PCBP1 in HCT116 PCBP1^WT/WT^, HCT116 PCBP1^L100Q/WT^ cells, HCT116 PCBP1^+/+^ and HCT116 PCBP1^+/-^ that were lysed in 8M Urea. Representative image is shown (n=3).

### PCBP1 L100/L102 mutants act in a dominant negative manner to suppress wild type PCBP1 expression

KH domain-containing RNA-binding proteins are known to associate with each other and form higher order structures such as dimers/tetramers^37,58–60^. To test whether PCBP1 wild type forms higher order structures and how this is affected by the L100Q mutation, we performed co-immunoprecipitation (IP) experiments. On HA immunoprecipitation, we detect binding between Flag-PCBP1 WT and HA-PCBP1 WT (Figure 6A, lane 4), indicating that PCBP1 wild type can bind to other wild type molecules. In this same HA IP, we observe an interaction between Flag-PCBP1 L100Q and HA-PCBP1 WT, indicating that wild type PCBP1 and mutant PCBP1 can associate (Figure 6A, lane 5). Similar results are obtained upon a reciprocal Flag-tag IP (Figure 6B).

**Figure 6:**
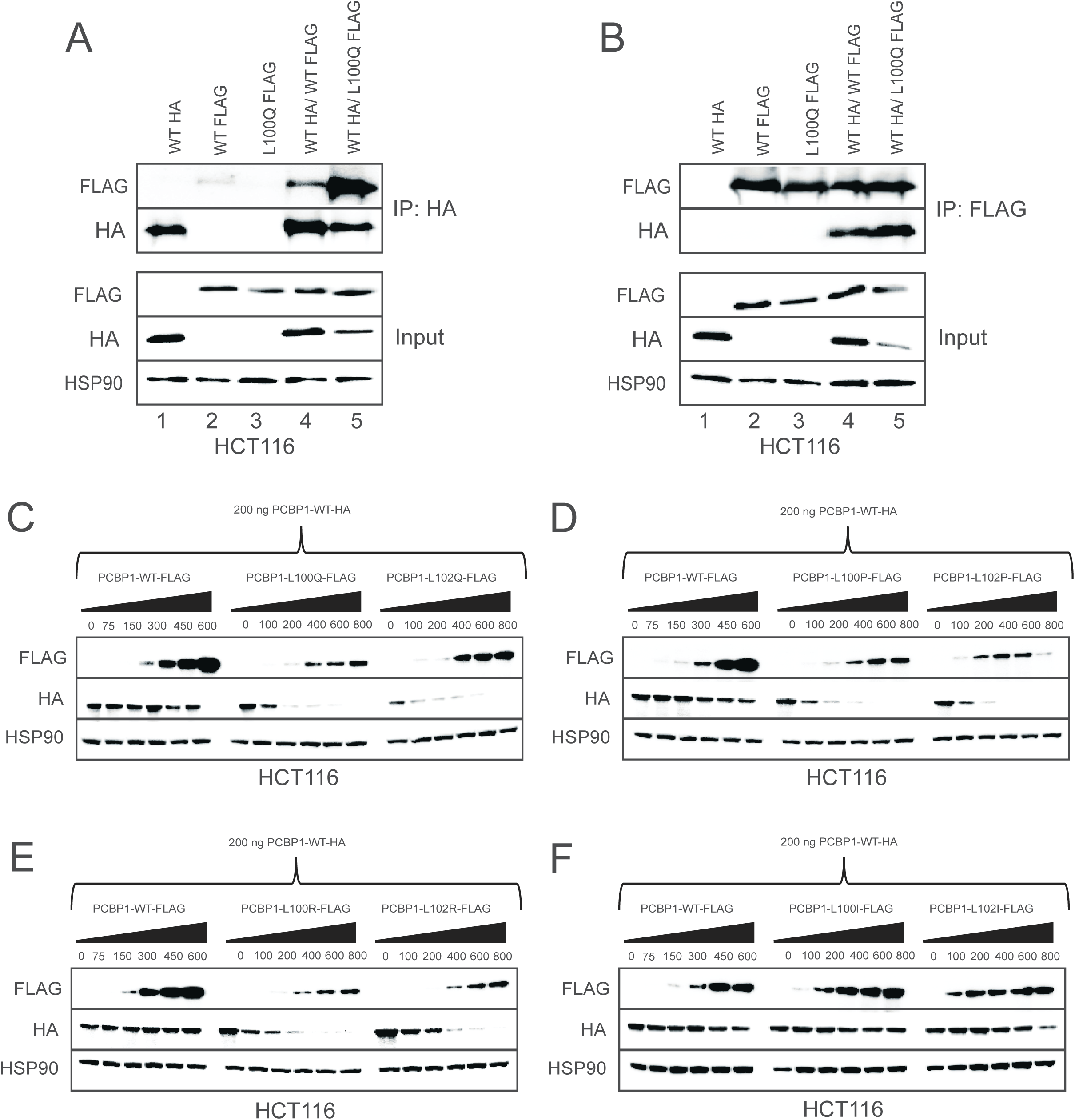
PCBP1 L100/L102 mutants act in a dominant negative manner to suppress wild type PCBP1 expression. **(A & B)** Co-immunoprecipitation experiments in HCT116 cells to interrogate binding between HA-tagged PCBP1 WT and Flag-tagged PCBP1 WT or L100Q using HA beads (A) or Flag beads (B). Representative image is shown (n=3). **(C-F)** Western blots to assess the dominant negative nature of the PCBP1 mutants. HCT116 cells were transiently transfected with a combination of 200 ng of HA-tagged PCBP1 WT and the indicated amounts of the Flag-tagged PCBP1 mutants. Representative image is shown (n=3).

Associations between wild type and mutant forms of tumor suppressor genes have been observed for NF1^33^, PTEN^34^, and TP53^63^, and in these cases the mutant protein typically acts in a dominant negative manner to impact the expression and/or function of the wild type protein. To assess whether PCBP1 L100/L102 mutations function via a dominant negative mechanism, we performed co-transfection experiments using HA-tagged PCBP1 WT with Flag-tagged PCBP1 WT or mutant plasmids in HCT116 cells. Each well of cells was transfected with 200 ng of HA-tagged PCBP1 WT along with increasing amounts (indicated in figure) of Flag-tagged PCBP1 WT or Flag-tagged PCBP1 L100/L102 mutants. The amount of PCBP1 L100/L102 mutant plasmids was adjusted based on Figure 4B so as to have a similar amount of mutant and wild type PCBP1 protein expression for these experiments. Here, we observe that increasing amounts of Flag-tagged PCBP1 WT have no effect on HA-tagged PCBP1 WT expression as indicated by unchanged levels of HA band (Figure 6C). Conversely, we observe that increasing amounts of Flag-tagged PCBP1 L100Q/L102Q cause a decrease in levels of HA-tagged PCBP1 WT as indicated by the decreasing levels of the HA band (Figure 6C). Similar results are seen for other cancer-associated PCBP1 mutations L100P/R and L102P/R (Figure 6D-E). To test whether this observed dominant negative effect is not an artifact of transfection, we used the PCBP1 L100I and L102I mutants and found that this isomeric mutation of leucine to isoleucine, which does not compromise PCBP1 protein stability, does not act in a dominant negative fashion to diminish PCBP1 WT expression levels (Figure 6F). This dominant negative nature of the PCBP1 L100/L102 mutants was observed in MC38 cells as well (Supplementary Figure 4A-D).

Given that KH domain-containing RBPs can not only self-associate but also associate with each other, we wanted to test whether PCBP1 L100/L102 mutations act in a dominant negative manner to suppress PCBP2 WT and hnRNPK WT expression. Here, we observe that PCBP1 L100Q mutations have a modest dominant negative effect on PCBP2 wild type expression (Supplementary Figure 4E) and hnRNPK wild type expression (Supplementary Figure 4F).

Since the L100/L102 residue is conserved in other KH domain containing RNA-binding proteins, we asked whether the cancer-associated PCBP2 L100R and hnRNPK L149M mutations are dominant negative as well. We observe that neither of these mutations are dominant negative and do not have any effect on the expression of wild type PCBP2 or wild type hnRNPK respectively (Supplementary Figure 4G-H).

Taken together, our data indicate that cancer-associated PCBP1 L100/L102 mutations are dominant negative and cause a decrease in expression of wild type PCBP1. These PCBP1 mutations have modest effect on other KH-domain containing RBPs like PCBP2 and hnRNPK. Disease-associated mutations in residues L100/L102 in PCBP2 and hnRNPK do not have dominant negative effects, indicating that this phenomenon is likely unique to PCBP1 biology.

## Discussion

Hotspot missense mutations in the tumor suppressor PCBP1 at residues L100 and L102 have been reported in colorectal cancers^24–28^. While it is known that PCBP1 mutations are correlated with poor clinical outcomes^31,32^, how PCBP1 L100/L102 mutations contribute to colorectal cancer initiation or progression remains largely unknown. Further, how these hotspot mutations impact PCBP1 expression and function has only recently been under investigation.

Our data shows that PCBP1 L100/L102 mutants are destabilized and have increased turnover. The expression of the mutants is partially rescued through the use of proteasomal degradation inhibitors. Both MG132 and Bortezomib inhibit the proteasomal function as well as the activity of some proteases. In the literature, PCBP1 has been reported to undergo both ubiquitination by the E3 ubiquitin ligase ARIH1^64^ and caspase-mediated proteolysis^65^. Whether the PCBP1 L100/L102 mutants are turned over through these mechanisms or a yet undiscovered novel mechanism remains to be determined. We can also increase the recovery of the PCBP1 L100/L102 mutants when using chaotropic protein isolation buffers containing urea to assist with the solubilization of mutant PCBP1. Our data indicate that the mutants are likely aggregation-prone, resulting in their insolubility and increased turnover. These findings are in line with Porter et al, who could not synthesize recombinant PCBP1 L100P protein as it aggregated out of solution^29^.

The KH1-KH2 domains of PCBP2 form an intramolecular pseudodimer^37^, and given the 94% sequence identity at the amino acid level for PCBP1 and PCBP2, we assume this dimerization also occurs in PCBP1. The L100/L102 mutations occur at the KH1-KH2 dimerization interface. Our molecular dynamics simulations show increased inter-domain movement for the PCBP1 L100/L102 mutants compared to wild type PCBP1, likely indicating the disruption of the KH1-KH2 binding. The NMR structure for the KH1-KH2 domains from Du et al show that this KH1-KH2 interface is stabilized by primarily hydrophobic interactions^37^. Therefore, one can rationalize how the mutation of hydrophobic leucine residues at positions 100 and 102 to bulkier, charged, or structure-breaking amino acids (glutamine, arginine, or proline), as observed in colorectal cancer, could disrupt the KH1-KH2 interface. The KH1-KH2 interface is also held together by hydrogen bonds between the backbone carbonyl groups and amide groups of Arg17/Val103 and Arg17/Arg101 and electrostatic interactions between the guanidino groups of Arg17/Arg101 and the ɛ-amino group of Lys160^37^, all of which are in close proximity to the L100/L102 mutation site.

Co-immunoprecipitation experiments show that wild type PCBP1 forms homo-oligomers and hetero-oligomers with wild type PCBP1 and the PCBP1 L100Q mutant, respectively. However, it is yet unknown whether there are different rate constants and binding affinities for the homo-oligomerization versus hetero-oligomerization process. Further, we do not know if PCBP1 forms dimers or other higher order structures and how this stoichiometry is disrupted in PCBP1^L100Q/WT^ cells.

Finally, our data shows that PCBP1 L100/L102 mutants suppress the expression of wild type PCBP1 in a dominant negative manner. Dominant negative mutations therefore represent a mechanism via which cancer cells severely downregulate tumor suppressor gene expression and/or function without needing to mutate all the alleles of the gene. Dominant negative effects are also the only way a cancer cell can severely reduce the expression of a tumor suppressor gene that is essential for cellular viability, such as in the case of PCBP1. Our data from cBioPortal showing that PCBP1 L100/L102 mutant alleles co-exist with wild type PCBP1 alleles support the dominant negative theory. We find that this dominant negative effect is only true for PCBP1 and not for other KH-domain containing RNA-binding proteins within the same family such as hnRNPK or PCBP2, indicating that this is likely a PCBP1-specific mechanism. This hypothesis is supported by tumor sequencing data wherein hotspot mutations at residues L100/L102 are more common in PCBP1 compared to PCBP2 or hnRNPK. Together, our data show that PCBP L100/L102 mutant proteins are destabilized, and they bind to and reduce the expression of wild type PCBP1 in a dominant negative manner (Figure 7).

**Figure 7:**
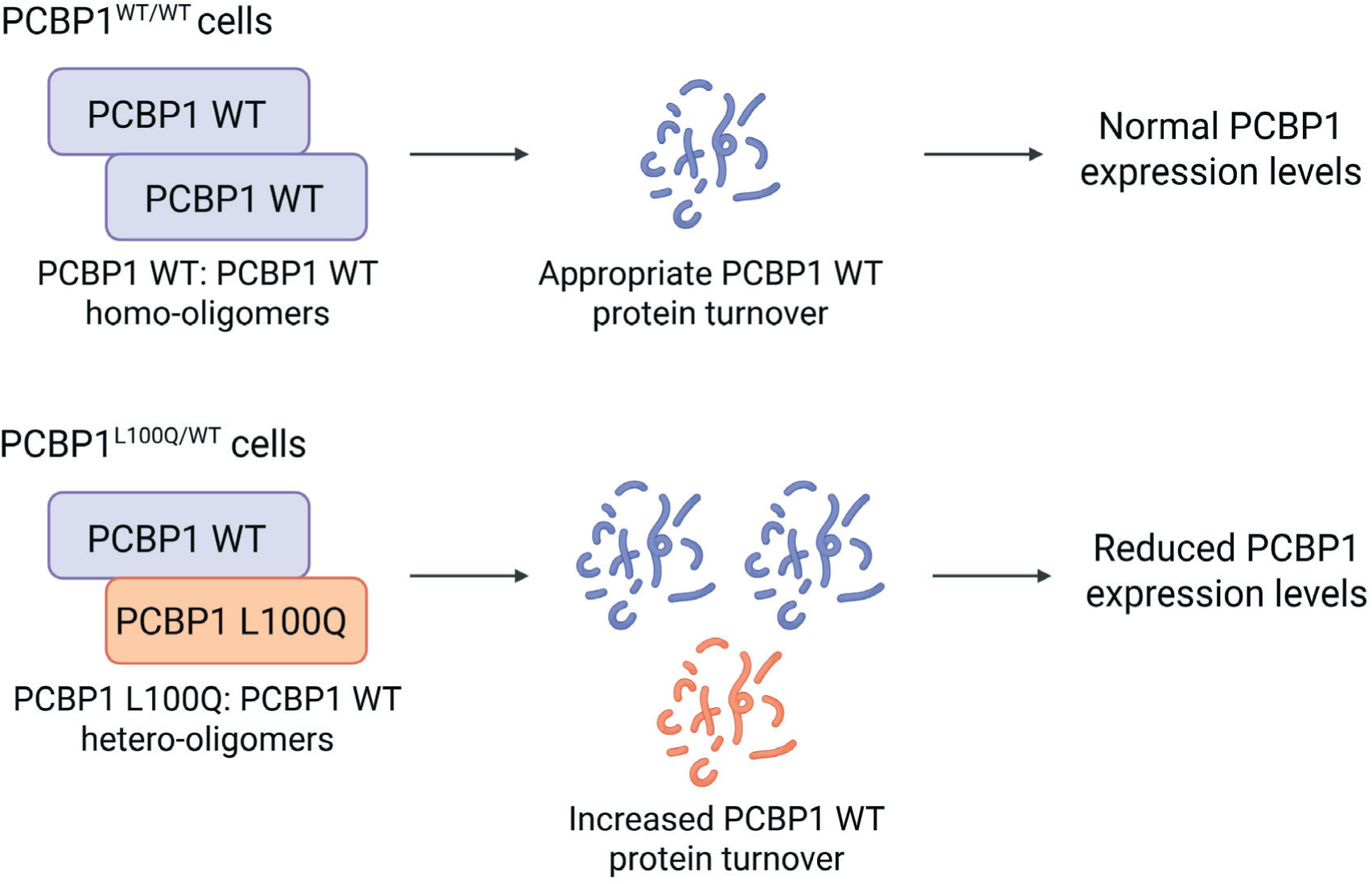
Model for mechanism of action of PCBP1 L100/L102 mutants. PCBP1 L100/L102 mutants are destabilized, have increased protein turnover and bind to and suppress the expression of wild type PCBP1 in a dominant negative manner.

There are several examples of dominant negative mutations observed for other tumor suppressor genes such as TP53, NF1, and PTEN. In case of TP53^63^ and NF1^61^, the mutant proteins are unstable and cause a loss of expression of total tumor suppressor gene expression. In contrast, dominant negative mutations in PTEN are more stable than the wild type PTEN^62^. However, these are catalytically dead, and upon heterodimerization with wild type PTEN, render it nonfunctional.

There are several questions that remain to be answered about these colorectal cancer associated PCBP1 mutations. One important consideration is how these mutations impact the function of PCBP1 as an RNA-binding protein. Data from Porter et al^29^ suggests that PCBP1 L100Q mutants can cross link three times more RNA than wild type PCBP1. However, they found no new RNA targets of PCBP1 L100Q when compared to wild type PCBP1. The observations of Porter et al allude to a gain of function phenotype for PCBP1 L100Q given its enhanced affinity for RNA. These seemingly disparate observations between Porter et al and our work can be contextualized by thinking about conformational changes in PCBP1. Tumor suppressor genes, like PTEN, are held in a closed, inactive conformation wherein their active sites are occluded to allow for cell growth. As needed, upstream signals cause a conformational change, which in the case of PTEN is dephosphorylation, that allows the protein to adopt a more open, active, albeit less stable, conformation^66^. It is plausible that PCBP1 L100/L102 mutants mimic the more open, active, but less stable conformation of PCBP1. Our molecular dynamic simulations showing a change in the association between the KH1 and KH2 domains in the PCBP1 mutants alludes to this conformational change. Data from Du et al show that the KH1-KH2 dimerization interface is on the opposite side of the RNA-binding interface, and it is very likely that the KH1-KH2 domains would have to dissociate from each other to allow for RNA binding. The PCBP1 L100/L102 mutants, with their impaired KH1-KH2 binding, may likely show enhanced RNA cross-linking due to the increased steric access to the RNA-binding interface. Some studies have reported that PCBP1 phosphorylation prevents its association with target mRNAs^67–69^, thereby inhibiting its function, akin to what is seen with PTEN. Detailed studies with scanning mutagenesis are needed to verify whether the PCBP1 L100/L102 mutants show the same conformational dynamics as observed with PTEN. Further, both Porter et al and our data show that the PCBP1 L100/L102 mutants are preferentially localized to the cytoplasm. Whether any changes in the function of the PCBP1 mutants are truly due to the mutation rather than a change in its sub-cellular compartmentalization is needed to understand how these mutations should be classified.

## Acknowledgments

We thank Dr. Kristina Godek and Dr. Duane Compton for use of the Ti2-E Inverted Microscope.

We thank Dr. Tyler Curiel for use of the QuantStudio 3 PCR machine. Flag-tagged PCBP1 WT and L100Q plasmids were a gift from Dr. Sean Post, MD Anderson Cancer Center.

## Contributions

PVB: experiment planning, execution and analysis, manuscript writing and editing

NCH: experiment planning, execution and analysis, manuscript writing and editing

MXA: experiment planning, execution and analysis, manuscript writing and editing

YL: experiment planning, execution and analysis

BB: experiment execution

UH: data analysis

EB: experimental planning, analysis and supervision

PM: experimental planning, analysis and supervision, manuscript writing and editing

