## Supplementary Materials for "Colorectal cancer-associated PCBP1 mutations disrupt protein stability in a dominant negative manner"

**Supplementary Figure Legends**

**Supplementary Figure 1: Mutations at L100/L102 disrupt PCBP1 protein expression and conformation. (A)** Western blot showing expression level for C-terminal Flag-tagged PCBP1 plasmids upon transient transfection in MC38 cells. Quantification of the western blot was done using three independent replicates and p-values were calculated using one-way ANOVA with a post hoc Dunnett's Test (** p<0.01) **(B)** Western blot showing expression level for C-terminal and N-terminal Flag-tagged PCBP1 plasmids upon transient transfection in HCT116 cells. **(C)** Heat map depicting AlphaMissense data to characterize the effect of different mutations on PCBP1 structure. **(D)** Relative root mean square fluctuation of the KH1-KH2 domains correlated with the expression levels of the various PCBP1 mutants in MC38 cells. RMSF across residues in the KH1 domain are calculated relative to residues in the KH2 domain. **(E and E')** Sub-cellular localization of PCBP1 mutants, as assessed by Flag immunofluorescence, upon transient transfection in HCT116 cells. Representative image is shown (scale = 50 μm; n=3).

**Supplementary Figure 2: Mutations at L100/L102 result in reduced expression of other KH domain containing RNA-binding proteins. (A & B)** Western blot showing expression level for C-terminal Flag-tagged PCBP2 and Flag-tagged hnRNPK plasmids respectively upon transient transfection in MC38 cells. Quantification of the western blot was done using three independent replicates and p-values were calculated using one-way ANOVA with a post hoc Dunnett's Test (* p<0.05; *** p<0.001; **** p<0.0001). **(C & D)** Western blot showing expression level for prostate cancer-associated C-terminal Flag-tagged PCBP2 L100R mutant in PC3 and DU145 cell lines respectively. Quantification of the western blot was done using three independent replicates and p-values were calculated using Mann Whitney test (ns p>0.05). **(E)** Western blot showing expression level for endometrial cancer-associated C-terminal Flag-tagged hnRNPK L149M mutant in HEC1A cells. Quantification of the western blot was done using three independent replicates and p-values were calculated using Mann Whitney test (* p<0.05).

**Supplementary Figure 3: The L100Q mutation destabilizes PCBP1 protein resulting in increased protein turnover. (A)** Copy number analysis for PCBP1 in HCT116 cells as assessed by droplet digital PCR. **(B)** *PCBP1* transcript levels in HCT116 $PCBP1^{WT/WT}$ and HCT116 $PCBP1^{L100Q/WT}$ cells as determined by qPCR. *PPIA* was used as a housekeeping control. Quantification was performed on three biological replicates and p-values were calculated using one-way ANOVA with a post hoc Dunnett's Test (** p<0.01; ns p>0.05). **(C)** Actinomycin chase assay to determine *PCBP1* mRNA stability in HCT116 $PCBP1^{WT/WT}$ and HCT116 $PCBP1^{L100Q/WT}$ cell lines. Quantification was performed on three biological replicates and p-values were calculated using two-way ANOVA with a post hoc Dunnett's Test (ns p>0.05). **(D&E)** Exponential decay curves for PCBP1 expression levels in HCT116 $PCBP1^{WT/WT}$ and HCT116 $PCBP1^{L100Q/WT}$ cells treated with cycloheximide alone (D) or cycloheximide + MG132 (E). **(F)** Cycloheximide chase assays with and without proteasomal inhibitor, bortezomib, in HCT116 $PCBP1^{WT/WT}$ and HCT116 $PCBP1^{L100Q/WT}$ cells. Representative image is shown (n=3)

**Supplementary Figure 4: PCBP1 L100/L102 mutants act in a dominant negative manner to suppress wild type PCBP1 expression. (A-D)** Western blots to assess the dominant negative nature of the PCBP1 mutants. MC38 cells were transiently transfected with a combination of 200 ng of HA-tagged PCBP1 WT and the indicated amounts of the Flag-tagged PCBP1 mutants. Representative image is shown (n=3). **(E-F)** Western blots to assess the dominant negative nature of the PCBP1 mutants on wild type PCBP2 and hnRNPK expression levels. HCT116 cells were transiently transfected with a combination of 200 ng of HA-tagged PCBP2 WT (E) or HA-tagged hnRNPK WT (F) and the indicated amounts of the Flag-tagged PCBP1 mutants. Representative image is shown (n=3). **(G)** Western blots to assess the dominant negative nature of the PCBP2 L100R mutant. PC3 cells were transiently transfected with a combination of 200 ng of HA-tagged PCBP2 WT and the indicated amounts of the Flag-tagged PCBP2 mutants. Representative image is shown (n=3). **(H)** Western blots to assess the dominant negative nature of the hnRNPK L149M mutant. HEC1A cells were transiently transfected with a combination of 200 ng of HA-tagged hnRNPK WT and the indicated amounts of the Flag-tagged hnRNPK mutants. Representative image is shown (n=3).

Supplementary Table 1

| **Primer Name** | **Sequence** |
|---|---|
| PCBP1 FL-F | ATGCCTCGAGCTCGCCATGGATGCCGGTGTGACTGAAA |
| PCBP1 FL-Flag-R | ATGCGAATTCCTACTTGTCATCGTCGTCCTTGTAGTCGCTGCACCCCATGCCCTT |
| PCBP1 FL-HA-R | ATGCGAATTCCTAAGCGTAATCTGGAACATCGTATGGGTAGCTGCACCCCATGCCCTT |
| PCBP1 L100Q-F | GGTCACCCAGAGGCTGGTG |
| PCBP1 L100Q-R | CACCAGCCTCTGGGTGACC |
| PCBP1 L100P-F | GGTCACCCCGAGGCTGGTG |
| PCBP1 L100P-R | CACCAGCCTCGGGGTGACC |
| PCBP1 L100R-F | GGTCACCCGGAGGCTGGTG |
| PCBP1 L100R-R | CACCAGCCTCCGGGTGACC |
| PCBP1 L100I-F | GGTCACCATCAGGCTGGTG |
| PCBP1 L100I-R | CACCAGCCTGATGGTGACC |
| PCBP1 L100V-F | GGTCACCGTGAGGCTGGTG |
| PCBP1 L100V-R | CACCAGCCTCACGGTGACC |
| PCBP1 L100M-F | GGTCACCATGAGGCTGGTG |
| PCBP1 L100M-R | CACCAGCCTCATGGTGACC |
| PCBP1 L100N-F | GGTCACCAACAGGCTGGTG |
| PCBP1 L100N-R | CACCAGCCTGTTGGTGACC |
| PCBP1 L102Q-F | CCCTGAGGCAGGTGGTGC |
| PCBP1 L102Q-R | GCACCACCTGCCTCAGGG |
| PCBP1 L102P-F | CACCCTGAGGCCGGTGGT |
| PCBP1 L102P-R | ACCACCGGCCTCAGGGTG |
| PCBP1 L102R-F | CACCCTGAGGCGGGTGGT |
| PCBP1 L102R-R | ACCACCCGCCTCAGGGTG |
| PCBP1 L102I-F | CCCTGAGGATCGTGGTGC |
| PCBP1 L102I-R | GCACCACGATCCTCAGGG |

| | |
|---|---|
| PCBP1 L102V-F | CCCTGAGGGTGGTGGTGC |
| PCBP1 L102V-R | GCACCACCACCCTCAGGG |
| PCBP1 L102M-F | CCCTGAGGATGGTGGTGC |
| PCBP1 L102M-R | GCACCACCATCCTCAGGG |
| PCBP1 L102N-F | CCCTGAGGAACGTGGTGC |
| PCBP1 L102N-R | GCACCACGTTCCTCAGGG |
| NFlag-PCBP1-F | ATGCCTCGAGCTCGCCATGGACTACAAGGACGACGATGACAAGGATGCCGGTGTGACTGAAA |
| NFlag-PCBP1-R | ATGCGAATTCCTAGCTGCACCCCATGCC |
| hnRNPK FL-F | ATGCCTCGAGATAAAAGAATATGGAAACTGAACAGCCAGAAGAAAC |
| hnRNPK FL-Flag-R | ATGCGAATTCTTACTTGTCATCGTCGTCCTTGTAGTCGAAAAACTTTCCAGAATACTGCTTCACACT |
| hnRNPK FL-HA-R | ATGCGAATTCTTAAGCGTAATCTGGAACATCGTATGGGTAGAAAAACTTTCCAGAATACTGCTTCACACT |
| hnRNPK L147Q-F | GACTGCGAGCAGAGGCTGTTG |
| hnRNPK L147Q-R | CAACAGCCTCTGCTCGCAGTC |
| hnRNPK L147P-F | CTGCGAGCCGAGGCTGTT |
| hnRNPK L147P-R | AACAGCCTCGGCTCGCAG |
| hnRNPK L147R-F | CTGCGAGCGGAGGCTGTT |
| hnRNPK L147R-R | AACAGCCTCCGCTCGCAG |
| hnRNPK L147I-F | GACTGCGAGATCAGGCTGTTG |
| hnRNPK L147I-R | CAACAGCCTGATCTCGCAGTC |
| hnRNPK L147V-F | GACTGCGAGGTGAGGCTGTTG |
| hnRNPK L147V-R | CAACAGCCTCACCTCGCAGTC |
| hnRNPK L147M-F | GACTGCGAGATGAGGCTGTTG |
| hnRNPK L147M-R | CAACAGCCTCATCTCGCAGTC |
| hnRNPK L147N-F | GACTGCGAGAACAGGCTGTTG |
| hnRNPK L147N-R | CAACAGCCTGTTCTCGCAGTC |
| hnRNPK L149Q-F | GAGTTGAGGCAGTTGATTCATCAG |
| hnRNPK L149Q-R | CTGATGAATCAACTGCCTCAACTC |
| hnRNPK L149P-F | TTGAGGCCGTTGATTCATCAG |
| hnRNPK L149P-R | CTGATGAATCAACGGCCTCAA |
| hnRNPK L149R-F | TTGAGGCGGTTGATTCATCAG |
| hnRNPK L149R-R | CTGATGAATCAACCGCCTCAA |
| hnRNPK L149I-F | TGCGAGTTGAGGATCTTGATTCATCAG |
| hnRNPK L149I-R | CTGATGAATCAAGATCCTCAACTCGCA |
| hnRNPK L149V-F | TGCGAGTTGAGGGTGTTGATTCATCAG |
| hnRNPK L149V-R | CTGATGAATCAACACCCTCAACTCGCA |
| hnRNPK L149M-F | TGCGAGTTGAGGATGTTGATTCATCAG |
| hnRNPK L149M-R | CTGATGAATCAACATCCTCAACTCGCA |
| hnRNPK L149N-F | TGCGAGTTGAGGAACTTGATTCATCAG |

| | |
|---|---|
| hnRNPK L149N-R | CTGATGAATCAAGTTCCTCAACTCGCA |
| PCBP2 FL - F | ATGCCTCGAGCTGCTCGACATGGACACCGGTGTGATTGAAG GT |
| PCBP2 FL FLAG – R | ATGCGAATTCCTACTTGTCATCGTCGTCCTTGTAGTCGCTGCT CCCCATGCCACC |
| PCBP2 FL HA - R | ATGCGAATTCCTAAGCGTAATCTGGAACATCGTATGGGTAGCT GCTCCCCATGCCACC |
| PCBP2 L100Q - F | CCGGTCACCCAGAGGCTGGTG |
| PCBP2 L100Q - R | CACCAGCCTCTGGGTGACCGG |
| PCBP2 L100P - F | CCGGTCACCCCGAGGCTG |
| PCBP2 L100P - R | CAGCCTCGGGGTGACCGG |
| PCBP2 L100R - F | CCGGTCACCCGGAGGCTG |
| PCBP2 L100R - R | CAGCCTCCGGGTGACCGG |
| PCBP2 L102Q - F | ACCCTGAGGCAGGTGGTC |
| PCBP2 L102Q - R | GACCACCTGCCTCAGGGT |
| PCBP2 L102P - F | ACCCTGAGGCCGGTGGTC |
| PCBP2 L102P - R | GACCACCGGCCTCAGGGT |
| PCBP2 L102R - F | ACCCTGAGGCGGGTGGTC |
| PCBP2 L102R - R | GACCACCCGCCTCAGGGT |

Supplementary Table 2

| Antibody | Supplier | Catalog Number | Experiment | Dilution |
|---|---|---|---|---|
| Anti-Flag M2 produced in mouse | Sigma | F1804-200UG | WB, IF | 1:1000 |
| Anti-Flag produced in rabbit | Sigma | F7425-.2MG | WB | 1:1000 |
| HA-Tag | Cell Signaling Technology | 3724 | WB | 1:1000 |
| Anti-PCBP1 | Abcam | AB168377 | WB | 1:1000 |
| Anti-HSP90 | Enzo | ADI-SPA-836-F | WB | 1:5000 |
| Anti-SOD1 | Cell Signaling Technlogy | 2770S | WB | 1:1000 |
| Mouse anti-rabbit IgG | Jackson Immuno Research | 211-032-171 | WB | 1:10,000 |
| Goat anti-mouse IgG | Jackson Immuno Research | 115-035-174 | WB | 1:10,000 |
| Invitrogen™ Donkey anti-Mouse IgG (H+L) Highly Cross-Adsorbed Secondary Antibody, Alexa Fluor™ 488 | Thermo Fisher Scientific | A21202 | IF | 1:1000 |
| Invitrogen™ Goat anti-Mouse IgG (H+L) Highly Cross-Adsorbed Secondary | Thermo Fisher Scientific | PIA32742 | IF | 1:1000 |

| | | | | |
|---|---|---|---|---|
| Antibody, Alexa Fluor™ Plus 594 | | | | |
| Invitrogen™ Goat anti-Rabbit IgG (H+L) Highly Cross-Adsorbed Secondary Antibody, Alexa Fluor™ Plus 488 | Thermo Fisher Scientific | PIA32731 | IF | 1:1000 |
| Invitrogen™ Goat anti-Rabbit IgG (H+L) Highly Cross-Adsorbed Secondary Antibody, Alexa Fluor™ 594 | Thermo Fisher Scientific | A11037 | IF | 1:1000 |
| DAPI | Sigma | D9542 | IF | 1:2000 |

Supp Fig 1

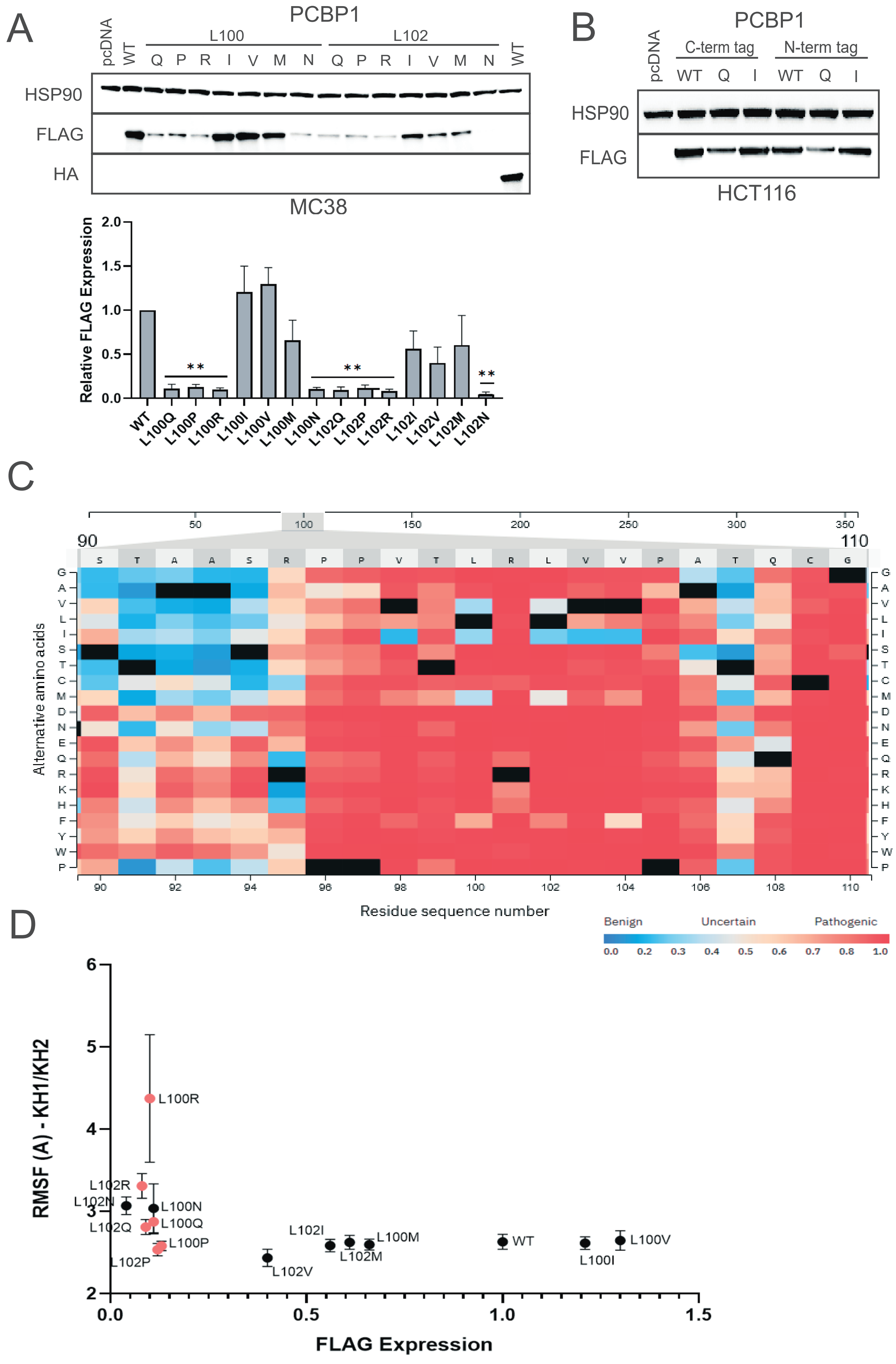
A
PCBP1
pcDNA
WT
L100
L102
Q P R I V M N Q P R I V M N
WT
HSP90
FLAG
HA
MC38
Relative FLAG Expression
2.0
1.5
1.0
0.5
0.0
**
**
**
WT
L100Q
L100P
L100R
L100I
L100V
L100M
L100N
L102Q
L102P
L102R
L102I
L102V
L102M
L102N
B
PCBP1
pcDNA
C-term tag
N-term tag
WT Q I WT Q I
HSP90
FLAG
HCT116
C
50
100
150
200
250
300
350
90
110
S T A A S R P P V T L R L V V P A T Q C G
G A V L I S T C M D N E Q R K H F Y W P
Alternative amino acids
90 92 94 96 98 100 102 104 106 108 110
Residue sequence number
Benign
Uncertain
Pathogenic
0.0 0.2 0.3 0.4 0.5 0.6 0.7 0.8 1.0
D
RMSF (A) - KH1/KH2
6
5
4
3
2
L100R
L102R
L102N
L100N
L102Q
L100Q
L100P
L102P
L102I
L102V
L100M
L102M
WT
L100I
L100V
0.0
0.5
1.0
1.5
FLAG Expression

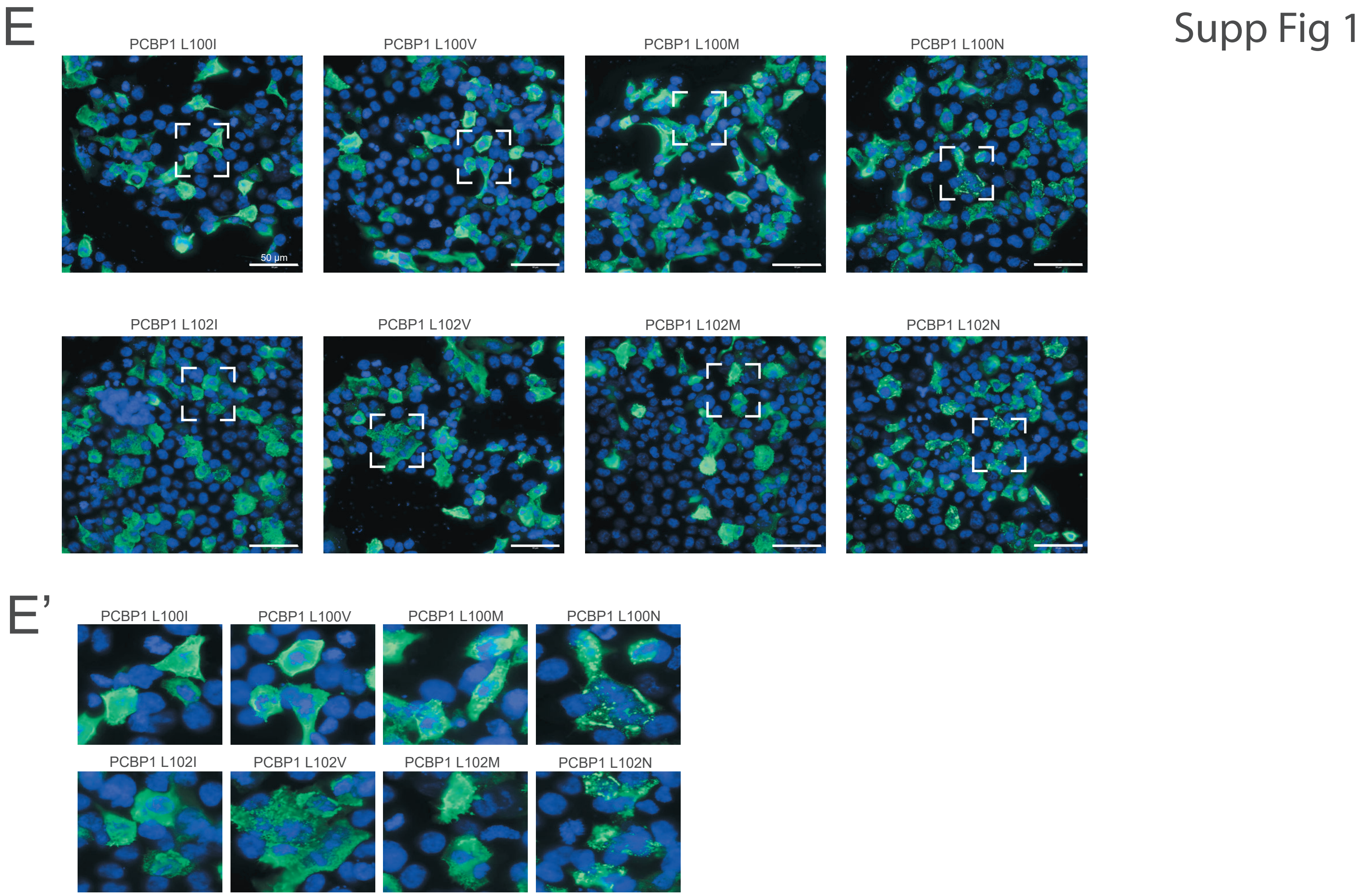
E
Supp Fig 1
PCBP1 L100I
PCBP1 L100V
PCBP1 L100M
PCBP1 L100N
50 µm
PCBP1 L102I
PCBP1 L102V
PCBP1 L102M
PCBP1 L102N
E’
PCBP1 L100I
PCBP1 L100V
PCBP1 L100M
PCBP1 L100N
PCBP1 L102I
PCBP1 L102V
PCBP1 L102M
PCBP1 L102N

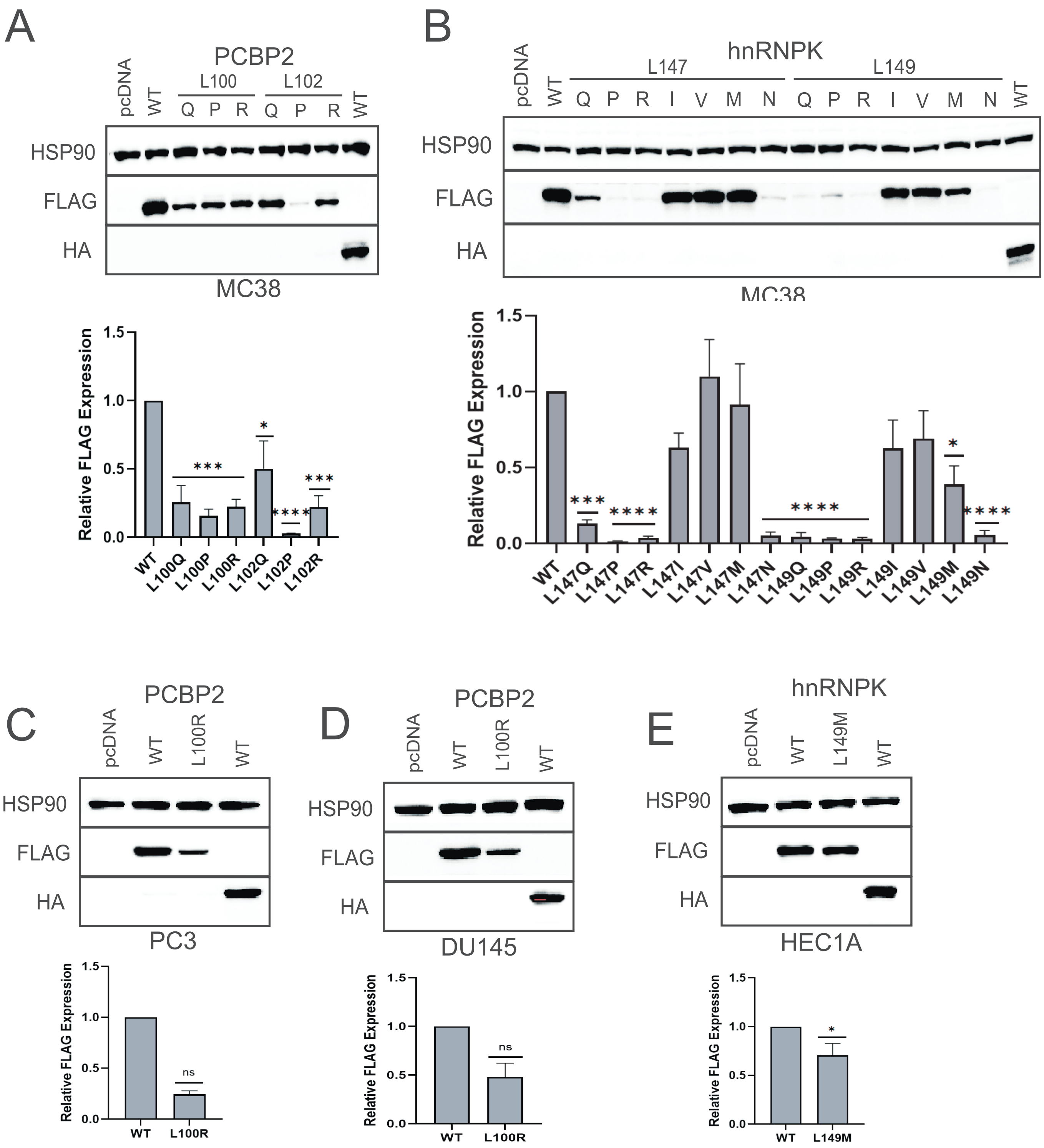
A
PCBP2
pcDNA
WT
L100
L102
Q
P
R
HSP90
FLAG
HA
MC38
Relative FLAG Expression
1.5
1.0
0.5
0.0
***
*
****
WT
L100Q
L100P
L100R
L102Q
L102P
L102R
B
hnRNPK
L147
L149
I
V
M
N
****
L147Q
L147P
L147R
L147I
L147V
L147M
L147N
L149Q
L149P
L149R
L149I
L149V
L149M
L149N
C
L100R
PC3
ns
D
DU145
E
L149M
HEC1A

Supp Fig 3

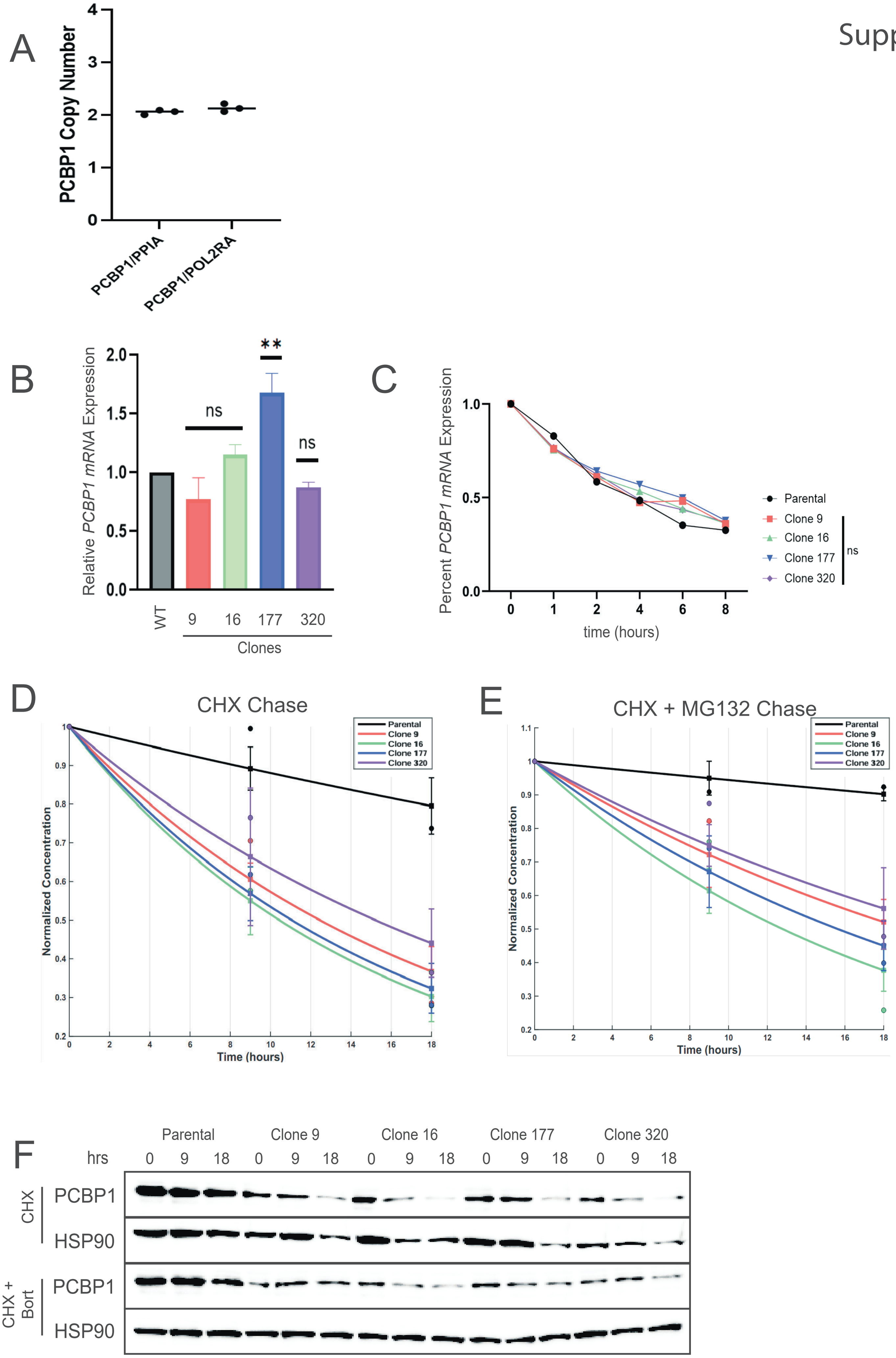
A
PCBP1 Copy Number
0
1
2
3
4
PCBP1/PPIA
PCBP1/POL2RA
B
Relative PCBP1 mRNA Expression
0.0
0.5
1.0
1.5
2.0
ns
**
ns
WT
9
16
177
320
Clones
C
Percent PCBP1 mRNA Expression
0.0
0.5
1.0
0
1
2
4
6
8
time (hours)
Parental
Clone 9
Clone 16
Clone 177
Clone 320
ns
D
CHX Chase
Normalized Concentration
Time (hours)
Parental
Clone 9
Clone 16
Clone 177
Clone 320
E
CHX + MG132 Chase
Normalized Concentration
Time (hours)
Parental
Clone 9
Clone 16
Clone 177
Clone 320
F
Parental
Clone 9
Clone 16
Clone 177
Clone 320
hrs
0 9 18 0 9 18 0 9 18 0 9 18 0 9 18
CHX
PCBP1
HSP90
CHX + Bort
PCBP1
HSP90

Supp Fig 4

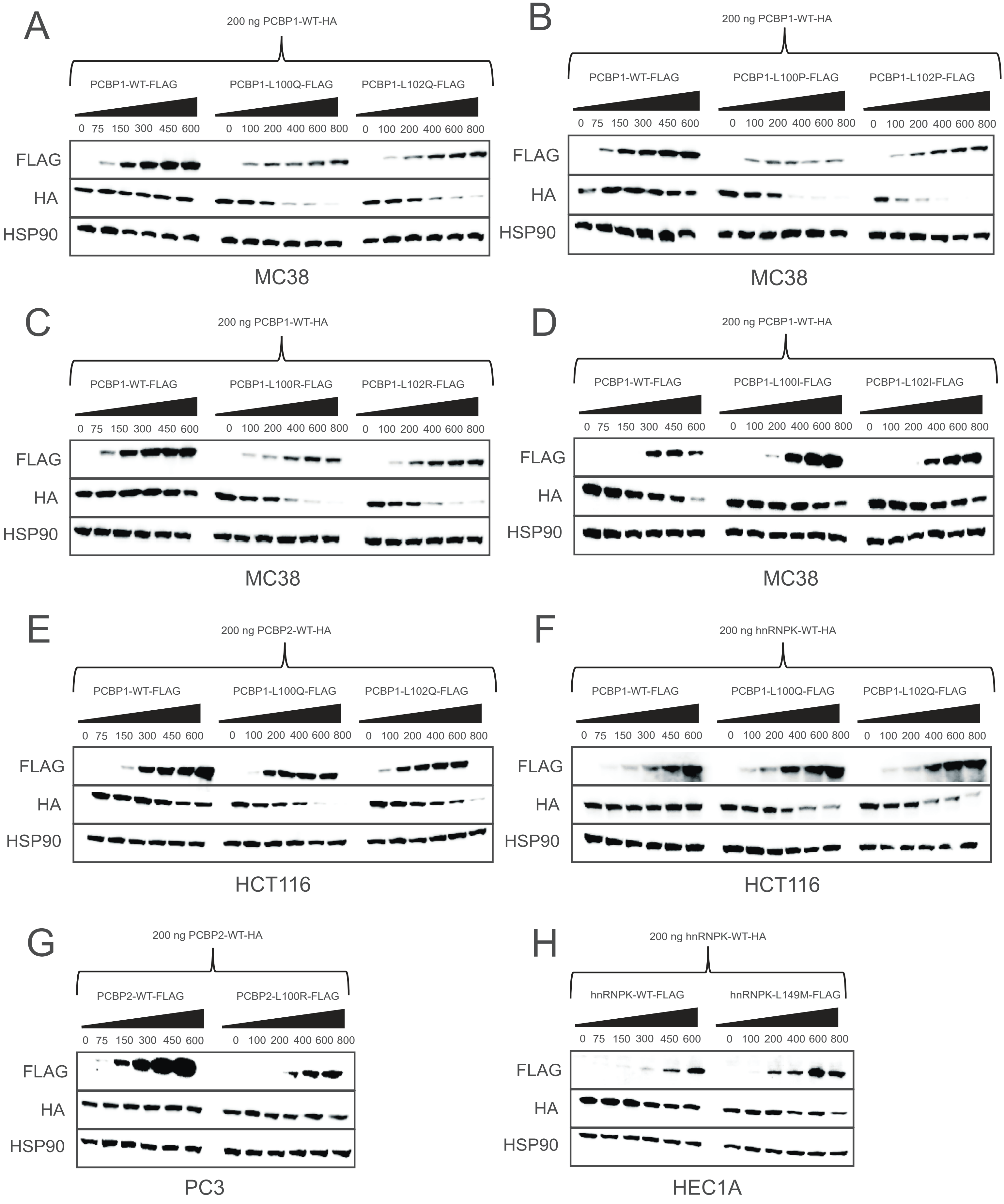
A
200 ng PCBP1-WT-HA
PCBP1-WT-FLAG
PCBP1-L100Q-FLAG
PCBP1-L102Q-FLAG
0 75 150 300 450 600
0 100 200 400 600 800
0 100 200 400 600 800
FLAG
HA
HSP90
MC38
B
200 ng PCBP1-WT-HA
PCBP1-WT-FLAG
PCBP1-L100P-FLAG
PCBP1-L102P-FLAG
0 75 150 300 450 600
0 100 200 400 600 800
0 100 200 400 600 800
FLAG
HA
HSP90
MC38
C
200 ng PCBP1-WT-HA
PCBP1-WT-FLAG
PCBP1-L100R-FLAG
PCBP1-L102R-FLAG
0 75 150 300 450 600
0 100 200 400 600 800
0 100 200 400 600 800
FLAG
HA
HSP90
MC38
D
200 ng PCBP1-WT-HA
PCBP1-WT-FLAG
PCBP1-L100I-FLAG
PCBP1-L102I-FLAG
0 75 150 300 450 600
0 100 200 400 600 800
0 100 200 400 600 800
FLAG
HA
HSP90
MC38
E
200 ng PCBP2-WT-HA
PCBP1-WT-FLAG
PCBP1-L100Q-FLAG
PCBP1-L102Q-FLAG
0 75 150 300 450 600
0 100 200 400 600 800
0 100 200 400 600 800
FLAG
HA
HSP90
HCT116
F
200 ng hnRNPK-WT-HA
PCBP1-WT-FLAG
PCBP1-L100Q-FLAG
PCBP1-L102Q-FLAG
0 75 150 300 450 600
0 100 200 400 600 800
0 100 200 400 600 800
FLAG
HA
HSP90
HCT116
G
200 ng PCBP2-WT-HA
PCBP2-WT-FLAG
PCBP2-L100R-FLAG
0 75 150 300 450 600
0 100 200 400 600 800
FLAG
HA
HSP90
PC3
H
200 ng hnRNPK-WT-HA
hnRNPK-WT-FLAG
hnRNPK-L149M-FLAG
0 75 150 300 450 600
0 100 200 400 600 800
FLAG
HA
HSP90
HEC1A
